# Molecular and cellular heterogeneity of gastric cancer explained by methylation-driven key regulators

**DOI:** 10.1101/2020.01.27.920744

**Authors:** Seungyeul Yoo, Quan Chen, Li Wang, Wenhui Wang, Ankur Chakravarthy, Rita Busuttil, Alex Boussioutas, Dan Liu, Junjun She, Tim R. Fenton, Jiangwen Zhang, Xiaodan Fan, Suet-Yi Leung, Jun Zhu

**Affiliations:** Icahn Institute for Data Science and Genomics Technology, Icahn School of Medicine at Mount Sinai, New York, NY, USA; Department of Genetics and Genomic Sciences, Icahn School of Medicine at Mount Sinai, New York, NY, USA; Sema4, a Mount Sinai venture, Stamford, CT, USA; Princess Margaret Cancer Centre, University of Toronto, Ontario, Canada; Department of Medicine, University of Melbourne, Parkville, Victoria, Australia; Department of Surgery, First Affiliate Hospital, Xi’an Jiaotong University, Xi’an, China; School of Biosciences, University of Kent, Canterbury, UK; School of Biological Sciences, University of Hong Kong, Hong Kong, China; Department of Statistics, Chinese University of Hong Kong, Hong Kong, China; Department of Pathology, University of Hong Kong, Hong Kong, China

## Abstract

Gastric cancer (GC) is a heterogeneous disease of diverse genetic, genomic, and epigenetic alterations. Tumor microenvironment (TME) also contributes to the heterogeneity of GC. To investigate GC heterogeneity, we developed an Integrative Sequential Causality Test (ISCT) to identify key regulators of GC by integrating DNA methylation, copy number variation, and transcriptomic data. Applying ISCT to three GC cohorts containing methylation, CNV and transcriptomic data, 11 common methylation-driven key regulators (*ADHFE1, CDO1, CRYAB, FSTL1, GPT, PKP3, PTPRCAP, RAB25, RHOH, SFN, and SORD*) were identified. Based on these 11 genes, gastric tumors were clustered into 3 clusters which were associated with known molecular subtypes, Lauren classification, tumor stage, and patient survival, suggesting significance of the methylation-driven key regulators in molecular and histological heterogeneity of GC. We further showed that chemotherapy benefit was different in the 3 GC clusters and varied depending on the tumor stage. Both immune/stromal proportions in TME and tumor cell genomic variations contributed to expression variations of the 11 methylation-driven key regulators and to the GC heterogeneity.

## Introduction

Gastric cancer (GC) is the fifth most common (8.2% of the total cancer cases) type of cancer and one of leading causes of global cancer death^1^. It occurs predominantly in Eastern Asian countries such as China, South Korea, and Japan^2^. In the United States, there were about 28,000 newly diagnosed GC patients and more than 10,000 deaths due to GC in 2017^3^. Many factors contribute to the high mortality and morbidity of GC. First, no distinct signs of symptoms of the disease until advanced stages and, hence, it is difficult to diagnose at early stage without routine screening via endoscopy^4^. Second, GC is heterogeneous in terms of biology, histology, and survival, and the diversity and complexity of the disease limits clear understanding of molecular mechanisms underlying in tumorigenesis, tumor progression, and metastasis of GC. Several studies have focused on the heterogeneity of GC in histology and/or in molecular characteristics. Lauren classification divides gastric tumors into three types (intestinal, diffuse or mixed-indeterminate) based on histology^5^. The diffuse type tumors are associated with poorer prognosis and develop at earlier age, while the intestinal gastric tumors are associated with intestinal metaplasia and advanced age^6, 7^. Recent studies identified distinct molecular alterations by gene expression, genetics, epigenetics, and proteomics in GC^8–11^. The Cancer Genome Atlas (TCGA) GC study categorizes GC into four subtypes 1) Epstein-Barr virus (EBV) with positive infection by the virus, 2) Microsatellite instability (MSI), 3) Chromosomal instability (CIN), and 4) Genome stable (GS) tumors based on molecular features such as viral infection, DNA methylation, genome stability, and mutation burden^10^.

DNA methylation alteration is one of key factors contributing to GC heterogeneity. EBV tumors exhibit excessive increases of DNA methylation in genome-wide scale, CpG island methylator phenotype (CIMP)^10, 12^. MSI GCs also show CIMP pattern while the level of hyper-methylation is lower than tumors in EBV group^10, 13^. GC clusters based on DNA methylation alterations such as CIMP were associated with clinical outcomes^9^. Also, it has been shown that genes involved in cancer-related pathways are more frequently affected by DNA methylations than by genetic alterations^14^. While these studies suggest a significant role of DNA methylation in gastric tumorigenesis and progression, detailed molecular mechanisms altered by DNA methylation changes are not well understood.

Copy number variation (CNV), a form of genomic alteration including amplification, or deletion of one or more sections of a chromosome, has a significant role in GC tumorigenesis^15, 16^. Previous studies^8, 17, 18^ identify multiple CNV regions in GC, including gains of 3p22, 4q25, 8q24, 11p13, and 20q13 and losses of 1p36 and 9p21. It has also been shown that the number of CNV occurrences is higher in patients with metastasis than in patients without metastasis^17^. Deng et al.^18^ identify 13 genomic amplifications and 9 deletions containing multiple cancer related genes such as *MYC*, *KRAS*, *CDK6* or *CDKN2A/B*. Wang et al.^8^ integrate multi-omics data and identify consistent chromosomal changes in previous studies including gains on chromosomes 1q, 5p, 7, 9, 12 and 20 and losses on 1p, 3p, 4, 5q, 9p, 17p, 18q, 19p, 21, and 22. While an increasing number of CNV regions have been identified in GC, it is still not fully understood how the associated genes impact tumorigenesis and progression of GC^19, 20^.

The histological, molecular, and genomic heterogeneity is further convoluted with heterogeneity in the tumor microenvironment (TME)^21, 22^. Cancer, immune, and stromal cells have distinct transcriptomic, genomic, and epigenetic patterns. In bulk tissue transcriptomic data, gene expression correlations may be due to TME heterogeneity or genomic heterogeneity (co-localized in deletion or amplification blocks) instead of transcriptional co-regulations, which hinder our understanding biology of GC. Also, the heterogeneity in TME is associated with prognosis or drug response. For example, higher proportion of immune cells in the tumor microenvironment is associated with better survival^23–25^ and better response to checkpoint blockade immunotherapies^26–28^. On the other hand, a higher proportion of stromal cells is associated with worse survival^29^, especially in GC^30–32^, and worse response to checkpoint blockade immunotherapies^33^. However, molecular connections of TME with different GC heterogeneities have not been systematically examined.

Here we describe an integrative causal model for identifying potential regulatory mechanisms in GC and their relationship with GC heterogeneity by integrating multi-omics data including gene expression, DNA methylation, and CNV profiles. Previously, we modeled a causality test as an empirical Bayesian estimation of the significance of *cis* DNA methylation regulated genes affecting other genes (*trans*)^34^. The previous causal model considered only one factor in gene expression regulation while there are multiple elements in the complex transcriptional regulation mechanisms. Therefore, we propose a new model here, Integrative Sequential Causality Test (ISCT), simultaneously considering transcriptional regulations driven by CNVs and methylation variations. We use the *cis*-association with promoter methylation and/or copy number alterations as priors and test whether the *trans*-associations are mediated through expression of *cis*-genes. (Figure 1A). Then, key regulators are inferred based on the number of downstream genes (Figure 1B). We applied ISCT to three GC cohorts (Supplementary Table 1) in which DNA methylation, CNV, and gene expression data were available, and highlighted key regulators and their downstream genes (Figure 1C). Eleven methylation-driven key regulators (*ADHFE1, CDO1, CRYAB, FSTL1, GPT, PKP3, PTPRCAP, RAB25, RHOH, SFN, and SORD*) were identified in all three datasets. Based on the common methylation-driven key regulators, GC samples were divided into three groups, each with distinct clinical outcome. The findings were validated in five independent GC cohorts (Supplementary Table 1). Among the 11 common methylation-driven key regulators, *FSTL1* expression was significantly associated with survival in 5 out of 7 datasets in which survival information were available. In GC cell lines, *FSTL1* was regulated epigenetically, and its correlated genes in cell lines were significantly enriched for its downstream genes inferred in bulk tissue profiles, consistent with a potential key regulator role of *FSTL1* in cell lines. To dissect the sources of gene expression variations of the 11 common methylation-driven key regulators, cellular compositions of bulk tissues were computationally inferred and their relationships with key regulators were investigated in the context of heterogeneity of gastric cancer (Figure 1C). *FSTL1* expression was highly correlated with the stromal cell fraction, suggesting both tumor intrinsic signals and variations in stromal cells in TME could regulate tumor cells.

**Figure 1.**
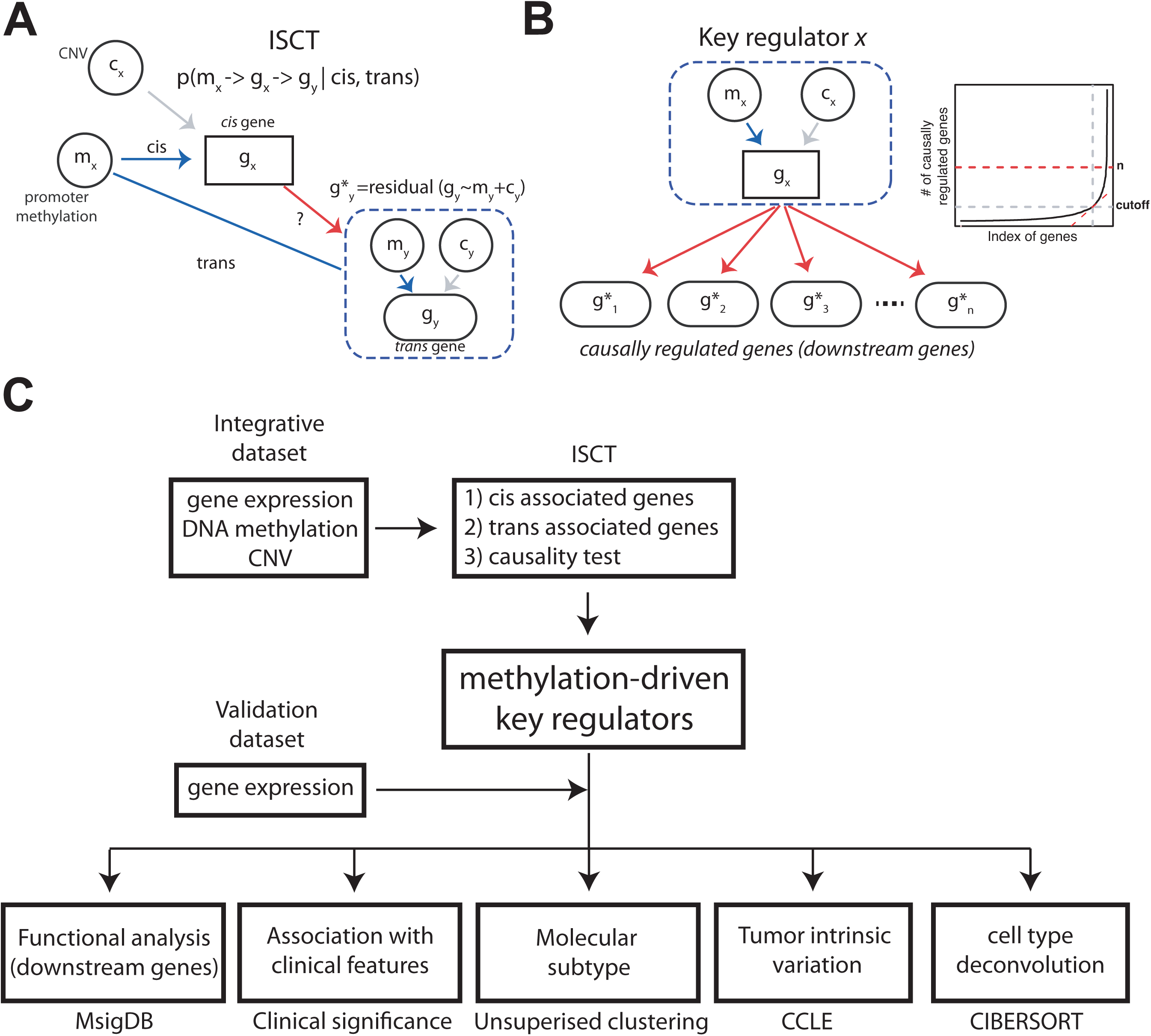
Overview of the study. **A.** ISCT: Given, *cis*- and *trans*-relationship among DNA methylation, CNV, and gene expression, the causal relationship between methylation status of gene *i* (*m*_*i*_) and expression of gene *j* (*g*_*i*_) is tested. The local methylation and CNV status of *trans* genes are also considered in the model using 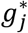 instead of *g*_*j*_. **B.** The definition of key regulator *x:* A *cis* gene *x* causally regulates significantly larger number of downstream genes above cutoff determined from a tangent line. **C.** Overall procedures of meta-analysis using ISCT and cell type deconvolution analysis from the three integrative datasets as well as the five gene-expression validation datasets.

## Results

### Comparison of ISCT and mediation tests on simulated datasets

Integrative Sequential Causality Test (ISCT) is developed for simultaneously identifying transcriptional regulations driven by CNVs and methylation variations. The ISCT approach and mediation tests^35, 36^ share some similarities in identifying relationships of gene expression *g* causally regulated by DNA methylation *m* and CNV *c* (Figure 1A). ISCT assumes sequential events occur in biology (Methods) while mediation tests take DNA methylation and CNV data as covariates without making any assumption. To systematically compare the performance of each method in identifying causal transcriptional regulations, we simulated two datasets, one based on a causal/mediation model and one based on an independent model according to the correlation distributions of the HKU dataset (detailed in Methods). We applied ISCT and two mediation tests to detect the underlying causal/mediation relationships. For the mediation tests, we conducted Baron and Kenny’s mediation test^36^ (referred to as “mediation test” in the following text and figures) to test for complete mediation, corresponding to the causal path tested in the ISCT approach; we also conducted the Sobel test^37^ to assess the significance of mediation effect, which does not necessarily guarantee complete mediation. In the dataset simulated based on a causal model (simulation #1), ISCT identified 91.7% of pairs as significant while the mediation test and the Sobel test identified 39.2% and 23.7% as significant, respectively (Figure 2A). In the dataset simulated based an independent model (simulation #2), the false positive rate of ISCT estimated by the proportion of simulated independent pairs tested significant was 0.46% while the false positive rates of the mediation test and the Sobel test were 0.40% and 0.31%, respectively (Supplementary Figure 1A). Therefore, the ISCT method demonstrated significantly higher power in detecting causal pairs comparing with other mediation tests while the false positive rate was well-controled under 0.5% similar to the other mediation tests.

**Figure 2.**
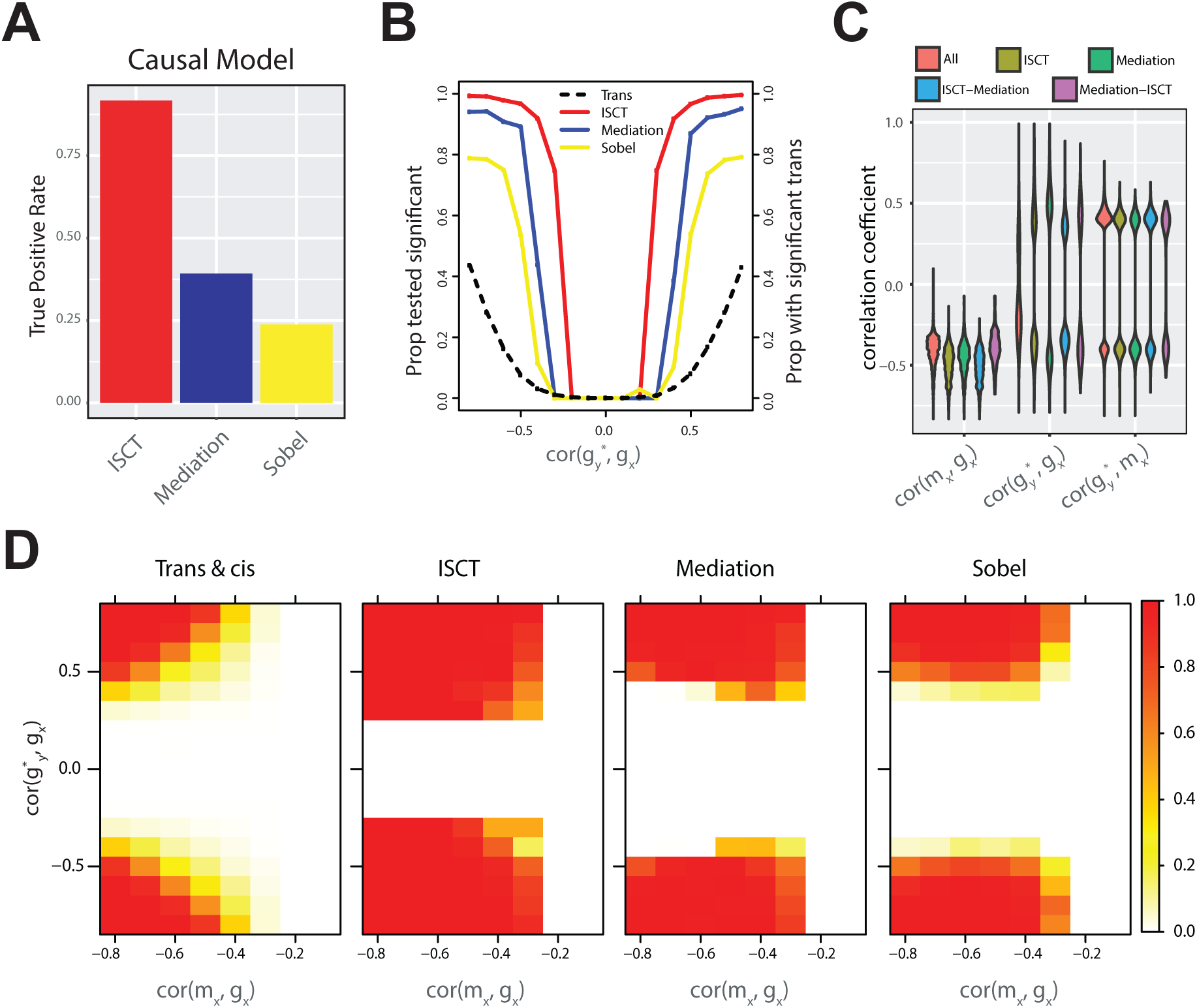
Simulation results to compare ISCT to mediation test. **A.** The true positive rate of each test estimated by the proportion of simulated causal pairs tested significant. **B.** For each pre-specified correlation level between *g*_*x*_ and 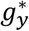, the dashed line shows the proportion of the simulated 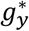’s showing significant *trans* relationship with *m_x_*; the yellow line shows the proportion detected by the Sobel mediation test with significant mediated relationship; the blue line shows the proportion detected by the mediation method with significant mediated relationship; the red line shows the proportion detected by the ISCT method with significant causal relationship. **C.** The pairwise correlation between *m*_*x*_, *g*_*x*_ and 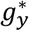 among 1) all the *cis*-*trans* gene pairs tested, 2) the significant pairs detected by the ISCT method, 3) the significant pairs detected by the mediation test, 4) the pairs detected by the ISCT method only, and 5) the pairs detected by the mediation test only. **D.** For each combination of a pre-specified *cor*(*g*_*x*_, *m*_*x*_) and *cor*(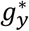, *g*_*x*_), the proportion of simulated *g*_*x*_, 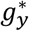 pairs with both significant *cis* and *trans* relationships with *m*_*x*_, and out of which the proportion detected by each method with significant causal/mediation relationship.

To systematically investigate the scenarios where these methods differ, we conducted further simulation studies given certain correlation levels among the variables to investigate the performance of each method under each specific scenario. We performed two additional simulations (simulations #3 and #4). In simulation #3, we randomly selected 10,000 *cis*-*trans* pairs from the HKU cohort with significant *cis*- and *trans*-relationships (detailed in Methods). This data-generating process mimiced the effect of a mediator^38^, by leaving the correlation between the dependent and the independent variables to vary. The power to identify causal relationships increased as the strength of the correlation between the dependent and the independent variables increased (Figure 2B). The ISCT method consistently outperformed the mediation test and the Sobel test with higher detection power at all correlation levels. At the correlations between the dependent and the independent variables range 0.3∼0.6 (Figure 2B), the Sobel test had the lowest detection power; the mediation test had minimal power at the correlation level of 0.3, around 40% power at the correlation level of 0.4, and reached over 85% power at the correlation level of 0.5; while ISCT had >75% power at the correlation level of 0.3, around 90% power at the correlation level of 0.4, and reached over 95% power at the correlation level of 0.5 (Figure 2B). These results suggested that the ISCT method was better powered in detecting causal pairs at lower correlation levels, where most pairs in the HKU cohort were observed (Figure 2C).

To investigate the performance of the two methods in the presence of colinearity, we simulated a fourth dataset (simulation #4, detailed in Methods). At the correlation between the dependent variable and the mediator around 0.5, the power of both mediation methods dropped with the increase in colinearity, and the drop in power with colinearity was more drastic when the correlation was around 0.4∼0.5 (Figure 2D). In contrast, the power of ISCT did not decrease with the increase in colinearity (Figure 2D). The results from simulations suggest that ISCT considering sequencial biological events in the model outperform typical mediation methods with higher detection power in the presence of colinearity.

### Methylation-driven key regulators

Both methylation and CNVs play a significant role in GC tumorigenesis and progression and should be simultaneously considered in transcriptional regulation as modeled in ISCT (Figure 1A).

#### 1) *cis* regulation driven by DNA methylation

We first investigated *cis* transcriptional regulations of *g*_*x*_ by DNA methylation in their promoter regions *m*_*x*_ and CNV alteration *c*_*x*_ based on a linear regression model, *g*_*x*_ ∼ *m*_*x*_ + *c*_*x*_ for gene *x* (detailed in Methods). At FDR<0.05, 7,917, 46,459, and 1,536 *cis*-methylation probes whose methylation levels were negatively correlated with the corresponding gene expression levels, which were summarized to 2,915, 11,880, and 1,119 *cis*-methylation genes, in the HKU, TCGA, and Singapore datasets, respectively. Among them, 474 *cis*-methylation genes were common among all three datasets (Figure 3A), which were enriched for signatures of ESTROGEN_RESPONSE_LATE and ESTROGEN_RESPONSE_EARLY in Hallmark gene sets from MSigDB (p=1.1×10^-11^ and 2.3×10^-5^, respectively). It is known that there is a higher GC risk in men than in women^39^ and estrogen plays an important role in GC tumorigenesis and progression^40^. We tested whether the methylation levels of the *cis*-methylation genes enriched for estrogen signatures were associated with the sex of patients, but this was not the case (Supplementary Figures 2A&B). In addition, most of them (17 and 24 out of 33 genes in HKU and Singapore dataset, respectively) were differentially methylated (t-test FDR<0.01) between tumor and adjacent normal tissues (Supplementary Figure 2C). Our result suggests that the estrogen response pathway is regulated at the epigenetic level.

**Figure 3.**
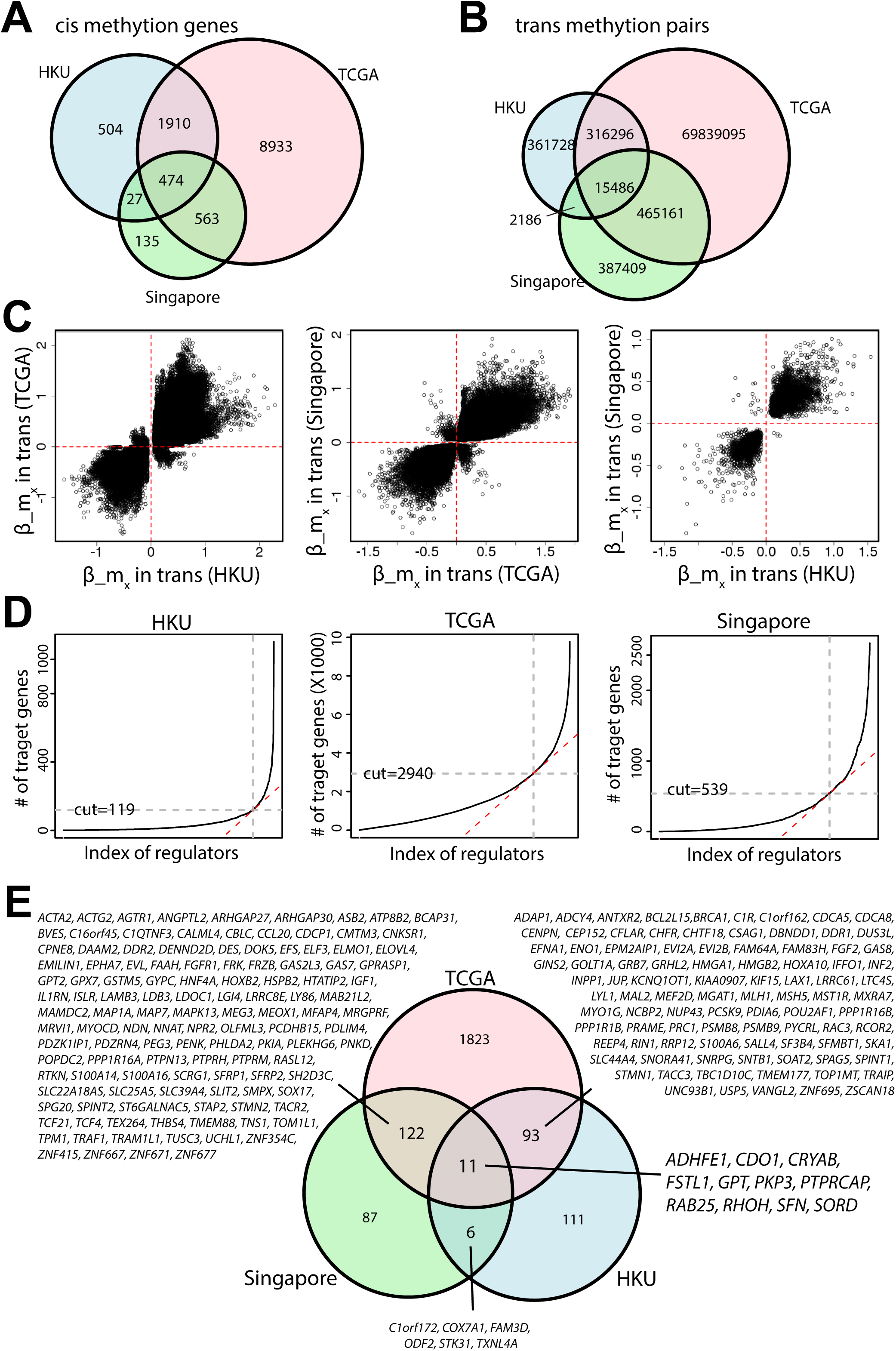
Identification of methylation-driven key regulators using ISCT. **A.** Comparison of the number of *cis* methylation genes identified from three GC cohorts. **B.** Comparison of the number of *trans* methylation pairs identified from three GC cohorts. **C.** Comparison of the directions of *trans* association of common *trans* genes among three GC cohorts. **D.** Identification of methylation-driven key regulators using cutoffs determined based on number of downstream genes in each GC cohorts. **E.** Common methylation-driven key regulators identified from three GC cohorts.

#### 2) *trans* regulations associated with DNA methylation

For each *cis*-methylation probe, we collected *trans*-methylation genes whose expression levels significantly correlated with the methylation variations (detailed in Methods). At the FDR<0.05, 695,696, 70,636,038, and 870,242 *trans*-methylation genes were identified in the HKU, TCGA, and Singapore datasets, respectively. The *trans*-methylation regulations significantly overlapped among different datasets (Figure 3B). Based on background of commonly significant *cis*-methylation genes among the three dataset, 70% and 66% of *trans* regulated gene pairs in HKU and Singapore were shared with the ones identified in the TCGA (FET Fold Enrichments (FEs)=2.3 and 2.1 with p<1.0×10^-256^ for HKU and Singapore, respectively). While with smaller percentage (about 5% and 18%), the *trans*–methylation gene pairs based on the HKU and Singapore dataset, respectively, were also significantly shared (FET FE=4.3 and p<1.0×10^-256^). Moreover, for the common *trans*-methylation gene pairs among different cohorts, the directions of the *trans* association were consistent (Figure 3C) suggesting similar impacts of methylation variations on the *trans* genes.

#### 3) Causal relationships driven by DNA methylation variations

After identifying *cis*- and *trans*-methylation genes, we further tested putative causal relationships between *cis*- and *trans*-methylation genes by testing whether a *cis*-methylation gene and a *trans*-methylation gene were independent given the expression of *cis*-genes (Methods). With the independence test p-value cutoff >0.01, 102,522, 19,085,499, and 353,576 causal gene pairs were identified in the HKU, TCGA and Singapore datasets, respectively (Supplementary Figure 3).

#### 4) Comparison of causal pairs based on ISCT and mediation tests

We took the HKU cohort as an example to investigate the difference between the causal relationships detected by ISCT and mediation test. Out of the 994,896 candidate *cis*-*trans* probe pairs, 143,332 pairs (summarized to 102,522 gene pairs) were identified as causal by ISCT while 53,267 pairs were identified as causal relationships with full mediation by Baron and Kenny’s mediation test. The majority of the causal pairs identified by Baron and Kenny’s mediation test were identified by the ISCT approach as well (Supplementary Figure 1B). To further investigate the scenarios where the two methods differ, we examined the distribution of gene expression correlations for the *cis*-*trans* gene pairs detected by each method. Both methods require a significant correlation between *g*_*x*_ and 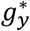 to detect a causal relationship, whereas the ISCT method was able to detect more pairs with significant causal relationships at a lower correlation level (Figure 2). In addition, the causal pairs detected by ISCT contained a higher faction of pairs with strong anti-correlations between *g*_*x*_ and *m*_*x*_, suggesting a possible power increase in the presence of colinearity.

#### 5) Methylation-driven key regulators

We next assessed whether there were any regulators whose methylation variations casually regulated expression levels of many other genes in *trans* (downstream genes), which were defined as methylation-driven key regulators (Methods). The number of downstream genes causally associated with each of *cis*-methylation genes followed a scale-free distribution (Supplementary Figure 4); most *cis*-methylation genes regulated a small number of downstream genes while a few *cis*-methylation genes regulated a large number of downstream genes. At the cutoffs based on reflection points (detailed in Methods, Figure 3D), 221, 2,049, and 226 methylation-driven key regulators were identified in the HKU, TCGA, and Singapore datasets, respectively. Among them, 232 were identified in at least two datasets and 11 were identified in all three datasets (Figure 3E and Supplementary Table 2). The 11 common methylation-driven key regulators were located on multiple chromosomes, chr1 (*RAB25* and *SFN*), chr3 (*FSTL1*), chr4 (*RHOH*), chr5 (*CDO1*), chr8 (*ADHFE1* and *GPT*), chr11 (*CRYAB*, *PKP3*, and *PTPRCAP*), and chr15 (*SORD*).

#### 6) Downstream genes of the methylation-based key regulators

For each key regulator, its downstream genes were further split into two groups; ones positively or negatively correlated with expression of the regulators. The downstream genes were significantly overlapped among all three datasets with consistent direction of regulations (Supplementary Table 2) suggesting similar impacts of the key regulators. We further determined downstream genes of the 11 common methylation-driven key regulators commonly identified at least in two datasets (Supplementary Table 3). Interestingly, the 11 methylation-driven key regulators shared similar set of their downstream genes (Figure EV1A). We performed functional enrichment test of the downstream genes of the methylation-driven key regulators against signature sets in MSigDB databases^41, 42^. The downstream genes of multiple methylation-driven key regulators were commonly enriched for three groups of Hallmark gene sets (Figure EV1B). One group of downstream genes, which were positively regulated by *FSTL1*, *CRYAB*, *CDO1*, and *ADHFE1* and negatively regulated by PKP3, SORD, GPT, RAB25, and SFN expression, was enriched for genes involved in EMT, MYOGENESIS (FET p-values=2.3×10^-15^ and 5.5×10^-14^, respectively for positively regulated downstream genes of *FSTL1*) while their anti-correlated downstream genes were enriched for cell cycle related functions such as E2F_TARGETS, G2M_CHECKPOINT, and MITOTIC_SPINDLE (FET p-values=4.8×10^-24^, 9.7×10^-14^ and 5.6×10^-6^, respectively for negatively regulated downstream genes of *FSTL1*). The downstream genes positively regulated by gene expression of *RHOH* and *PTPRCAP* were enriched for immune related functions such as INFLAMMATORY_RESPONSE, ALLOGRAFT_REJECTION, and INTERFERON_GAMMA_RESPONSE (for positively regulated downstream genes of *RHOH* p-values=5.2×10^-18^, 3.2×10^-37^, and 1.5×10^-16^, respectively).

### GC subtypes based on methylation-driven key regulators

#### Methylation-driven key regulators were correlated in GC

Based on gene expression similarity, the 11 common methylation-driven key regulators were clustered into two anti-correlated groups with *CDO1, CRYAB, FSTL1, ADHFE1, RHOH,* and *PTPRCAP* in one group and *GPT, SORD, PKP3, RAB25* and *SFN* in the other (Figure 4A). The first group was further separated into two gene clusters in which *CDO1, CRYAB*, *FSTL1,* and *ADHFE1* (G1) were co-expressed while *RHOH* and *PTPRCAP* (G2) were closely expressed. Five genes in the other group, *GPT, SORD, PKP3, RAB25,* and *SFN* (G3), were also highly correlated with one another. These patterns were consistently observed across all GC cohorts in the study (Figure 4A).

**Figure 4.**
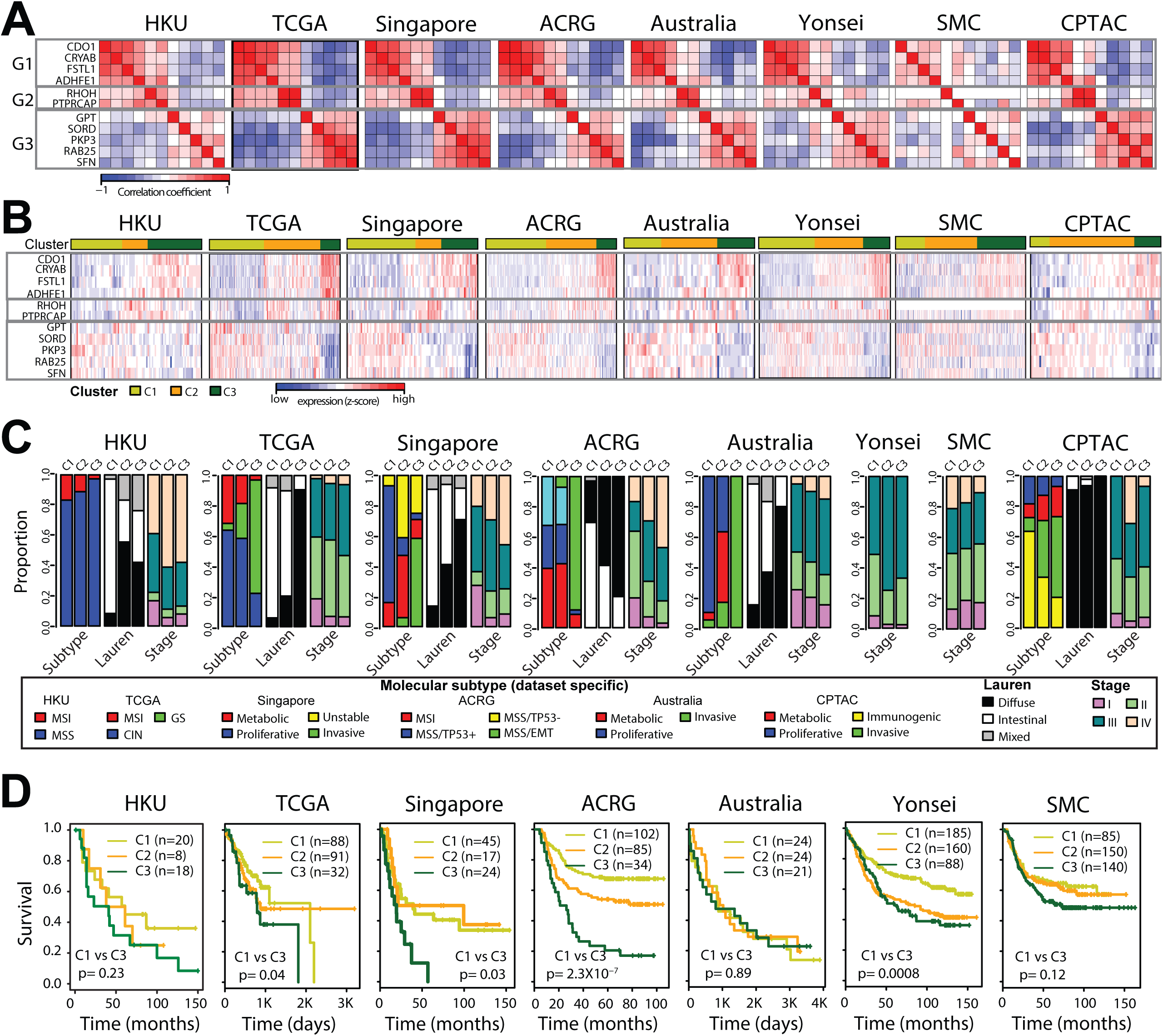
Tumor clusters based on the gene expression levels of the 11 methylation-driven key regulators. **A.** Co-expression of the 11 methylation-driven key regulators in all GC cohorts was measured as Pearson correlation coefficients. *CDO1*, *CRYAB*, *FSTL1*, and *ADHFE1* were clustered in one group (G1), *RHOH* and *PTPRCAP* in another (G2) and the rest (*GPT*, *SORD*, *PKP3*, *RAB25*, and *SFN*) in the other group (G3). In the SMC datasets, *RHOH* was not profiled and NA values were colored in white. **B.** K-mean clustering of GC tumors (k=3) based on expression levels of the methylation-driven key regulators. **C.** The proportion of molecular subtypes, Lauren class, and tumor stages within the tumor clusters. The molecular subtypes were determined in the original study of each cohort. For the Yonsei and SMC datasets, only stage information was available. **D.** KM-plots for survival analysis (Overall survival) among the tumor clusters. The number of samples in each cluster is shown. Survival differences between C1 and C3 were measured as likelihood ratio test p-values. For the HKU datasets, patients with palliative treatments were removed for survival analysis.

#### Tumor clusters based on the gene expression of methylation-driven key regulators

Tumor samples were clustered by k-means clustering into three groups based on the similarity of the expression of the 11 common methylation-driven key regulators (Figure 4B). The three groups, C1 to C3, showed increasing expression levels of genes in G1 and G2, while decreasing expression of genes in G3 and the samples in all GC datasets were clustered in a similar manner (Figure 4B). Samples in the groups C1 and C3 showed opposite expression patterns for the genes in G1 and G3. C2 samples showed intermediate expression of genes in G1 and G3 (Figure 4B).

Then, we investigated whether the tumor clusters based on key methyl regulator expression levels were associated with previously known molecular features of GC. The tumor clusters were significantly overlapped with Lauren classifications (Figure 4C and Supplementary Table 4). The C1 cluster was enriched for the intestinal subtype in all datasets except for CPTAC dataset in which most of samples are the diffuse subtype (FET FEs= 2.77, 1.5, 2.07, 1.97, and 1.99 and p-values= 4.5×10^-8^, 1.1×10^-5^, 0.0003, 2.8×10^-7^, and 0.005 in HKU, TCGA, Singapore ACRG, and Australia datasets, respectively). On the other hands, the C3 group was enriched for diffuse subtype (FET FEs= 1.75, 6.01, 3.14, 1.93, and 2.99 and p-values= 0.05, 6.0×10^-17^, 4.9×10^-5^, 3.3×10^-5^, and 3.9×10^-5^ in HKU, TCGA, Singapore, ACRG, and Australia datasets, respectively).

In addition, tumors in C1 significantly overlapped with the MSI and CIN subtype in TCGA (FET FEs=2.17 and 1.3, p-values=0.003 and 0.02, respectively), the proliferative subtype in Singapore and Australia datasets (FET FEs=7.74 and 4.69, p-values=2.8×10^-10^ and 1.4×10^-7^, respectively), and also the MSS/TP53-subtype in ACRG (FET FE=1.83, p-value=0.009), and Immunogenic subtypes in CPTAC dataset (FET FE=2.11 with p-value=0.04). Tumors in C2 significantly overlapped with the metabolic subtype in Singapore and Australia datasets (FET FEs=2.58, 12.19, p-values=0.03, 4.5×10^-6^, respectively). Tumors in the C3 were enriched for GS subtype in the TCGA dataset (FET FE=5.37, p-value=8.6×10^-12^). Interestingly, the C3 cluster was also highly enriched for mesenchymal-like subtype tumors in Singapore, ACRG, and Australia (FET FEs=18.08, 27.5, and 9.6, p-values=3.9×10^-8^, 7.8×10^-27^ and 2.4×10^-13^, respectively). While a previous study reported the TCGA GS subtype was not equivalent to the ACRG MSS/EMT subtype^43, 44^, our results suggest that they shared similar molecular features characterized by the 11 methylation-driven key regulators.

The C1-3 clusters were also associated with tumor stages in TCGA, Singapore, ACRG, and Yonsei datasets (Figure 4C and Supplementary Table 4). The results were consistent with previous reports that mesenchymal-like GC tumors were associated with diffuse type and more advanced tumors while the proliferative subtype was associated with intestinal type^10, 43, 45^. Taken all together, these results further suggest significant roles of the methylation-driven key regulators in the heterogeneity of GC tumors.

#### Survival differences among the tumor clusters

The clusters based on the methylation-driven key regulators showed prognostic differences (Figure 4D). In general, patients in the C1 group had better survival than ones in the C3 and the C2 group had intermediate survival, whether more closely to C1 or C3 depending on their similarity of expression patterns of the methylation-driven key regulators (Figure 4B). The survival differences between tumors in the C1 and C3 were statistically significant in TCGA, Singapore, ACRG, and Yonsei datasets (Likelihood Ratio (LR) test p-values=0.04, 0.03, 2.3×10^-7^, and 0.0008, respectively). In HKU and SMC datasets, the C1 and C3 clusters showed different survival compared to each other, while LR test p-values were not significant. The clusters were not associated with patients’ survival in the Australia datasets. It was worthy to note that the patients in Australia dataset were mixed with or without 5-FU treatment which showed significant survival differences within the same subtypes^45^. Combining all results above, the methylation-driven key regulators classify samples into three groups, which well agreed with molecular and histological heterogeneity as well as differences in clinical outcomes of gastric cancer.

#### Chemotherapy response differences among tumor clusters

A recent study reported by Oh et al. showed distinct genomic features between epithelial and mesenchymal phenotype (EP and MP) of gastric tumors^46^. They used a set of 299 signature genes, which included *FSTL1* and *GPT,* to classify gastric tumors into two groups. When comparing the tumor clusters based on the 11 methylation-driven key regulators with their EP/MP results using overlapping samples in the TCGA and ACRG datasets, C1 and C3 almost exclusively overlapped with EP and MP, respectively (Figures 5A, details in Appendix, Supplementary Table 5). This is consistent with the result that the positively regulated downstream genes of *FSTL1* were enriched for EMT pathways (FET p-values=2.3×10^-15^, EV1B). We analyzed the 4 additional GC datasets used in Oh et al study (Methods), and tumor clusters based on the 11 methylation-driven key regulators showed similar associations with EP and MP groups (Figures 5B&C, Supplementary Table 5). These results are consistent with our previous observation that C3 tumors were significantly associated with invasive or mesenchymal subtype of tumors in the Singapore and Australia datasets (Figure 4C and Supplementary Table 4). The tumors in C1 and C3 groups showed significant differences in overall survival as well as recurrence free survival (Figures 5D&E) consistently with distinct survival patterns between EP and MP groups^46^. These results suggest that the epithelial/mesenchymal phenotype can be largely explained by our methylation-driven key regulators.

**Figure 5.**
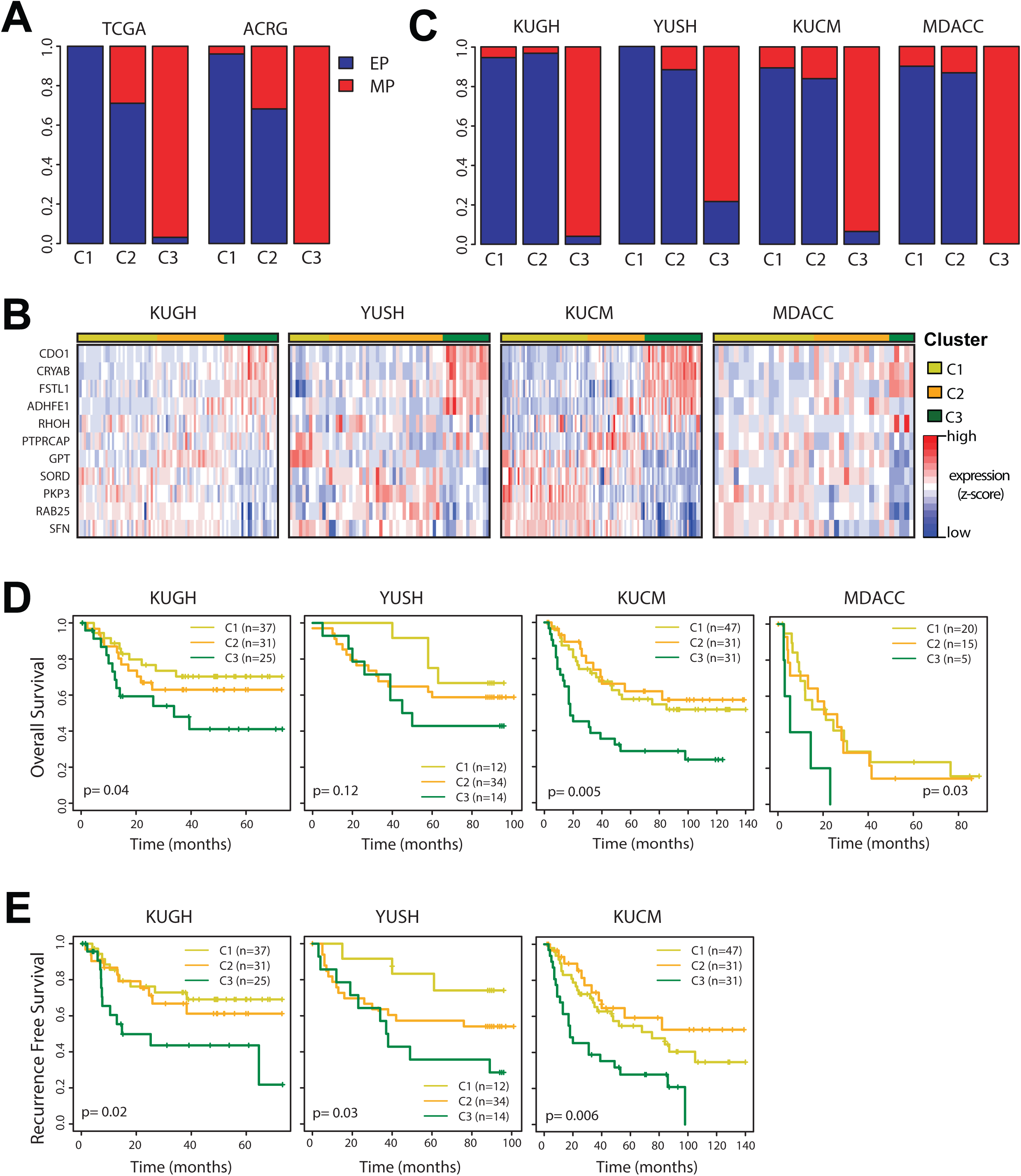
Tumor clusters based the 11 methylation-driven key regulators overlap with epithelial/mesenchymal phenotypes. **A.** Comparison of the tumor clusters based on the 11 methylation-driven key regulators (Figure 4B) with Epithelial/Mesenchymal subtypes determined in Oh et al.’s report^46^. Barplot showing overlapping rate (Supplementary Table 5) between two clustering results in TCGA and ACRG datasets. **B.** K-mean clustering of GC tumors (k=3) based on expression levels of the methylation-driven key regulators for 4 datasets not included in our study (KUGH, YUSH, KUCM, and MDACC). **C.** Comparison of the tumor clusters from Figure 7B with Epithelial/Mesenchymal subtypes determined in Oh et al.’s report^46^. **D & E.** Kaplan-Meier plots showing overall survival (D) and recurrence free survival (E) of each cluster. P-values indicate the significance of survivals between C1 and C3. Recurrence free survival for MDACC was not available.KM-plots for survival analysis (Overall survival) among the tumor clusters. The number of samples in each cluster is shown. Survival differences between C1 and C3 were measured as likelihood ratio test p-values.

Oh et al reported that adjuvant chemotherapy (CTX) was effective exclusively for EP tumors^46^. Following their procedure, we combined patients from KUGH, YUSH, and KUCM datasets, observed differences in the recurrence free survival between patients with and without CTX were significant only for C1 group (p=0.039) but not for C2 and C3 groups (p=0.44 and 0.12, respectively, Figure 6A), and similarly significantly in the EP cluster not in MP cluster (p=0.009 and 0.98, respectively, Figure 6A) as reported by Oh et al.

**Figure 6.**
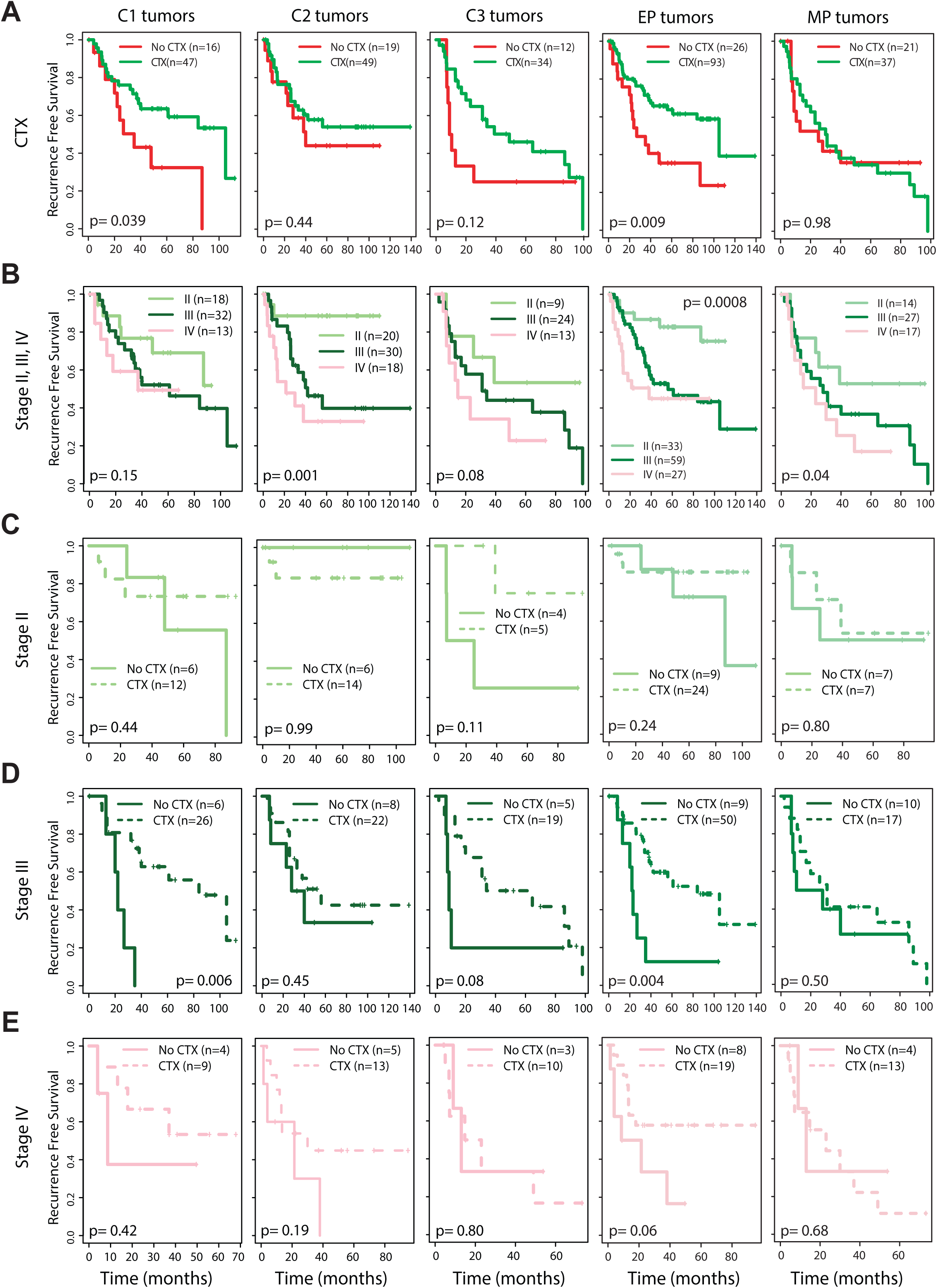
Adjuvant chemotherapy (CTX) sensitivity depending on tumor subtypes as well as tumor stages. **A.** Survival differences between patients with and without CTX in each group. C1-C3 tumors based on our clustering method and EP and MP subtypes reported by Oh et al.^46^ **B.** Survival differences among different tumor stages (II, III, and IV) in each group. **C.** Association between CTX and progression free survival at stage II. **D.** Association between CTX and progression free survival at stage III. **E.** Association between CTX and progression free survival at stage IV.

However, in each tumor cluster (C1-3 clusters by our method or EP/MP cluster by Oh et al), there was a mixture of tumors at different stages with poor survival for patients of advanced stage tumors (Figure 6B). When examining tumors according to molecular clusters and stages, patients benefited from CTX only when tumors were in the C1 cluster and at Stage III (p=0.006) based on our clustering method or in the EP cluster and at Stage III (p=0.004) based on Oh et al (Figures 6C-E). There might be CTX treatment benefit for tumors in the C3 cluster and at Stage II (p=0.11) or at stage III (p=0.08). These results suggest strong needs for personalizing treatment decision based on molecular phonotype as well as tumor stage.

### Clinical significance of the methylation-driven key regulators

Next, we associated the expression levels of the methylation-driven key regulators in tumors with patients’ survival and tumor stages in the 7 datasets (3 integrative datasets and 4 gene expression validation datasets except for CPTAC) and strong associations between them were observed with consistent directions among multiple datasets (Figures 7A&B, Supplementary Table 6). The increased expressions of the G1 genes (*CDO1*, *CRYAB*, *FSTL1*, and *ADHFE1*) were significantly associated with poor prognosis as well as advanced tumor stages in multiple datasets. A recent study showed that *CRYAB* overexpression induced invasion and migration via EMT in gastric cancer cells^47^. *FSTL1* expression showed the most consistent and strongest association with both survival and tumor stages within 5 datasets (survival: p-values=0.0002, 1.1×10^-6^, 0.02, 0.0004, and 0.04 for Singapore, ACRG, Australia, Yonsei and SMC datasets, respectively; tumor stages: p-values= 0.0003, 0.01, 6.9×10^-5^, 0.002, and 9.3×10^-5^ for HKU, TCGA, Singapore, ACRG, and Yonsei datasets, respectively). *FSTL1* was reported as a key mediator in immune dysfunction driven by metastasis and aging in mouse cancer models^48^ but no functions of it in gastric cancer was reported previously.

**Figure 7.**
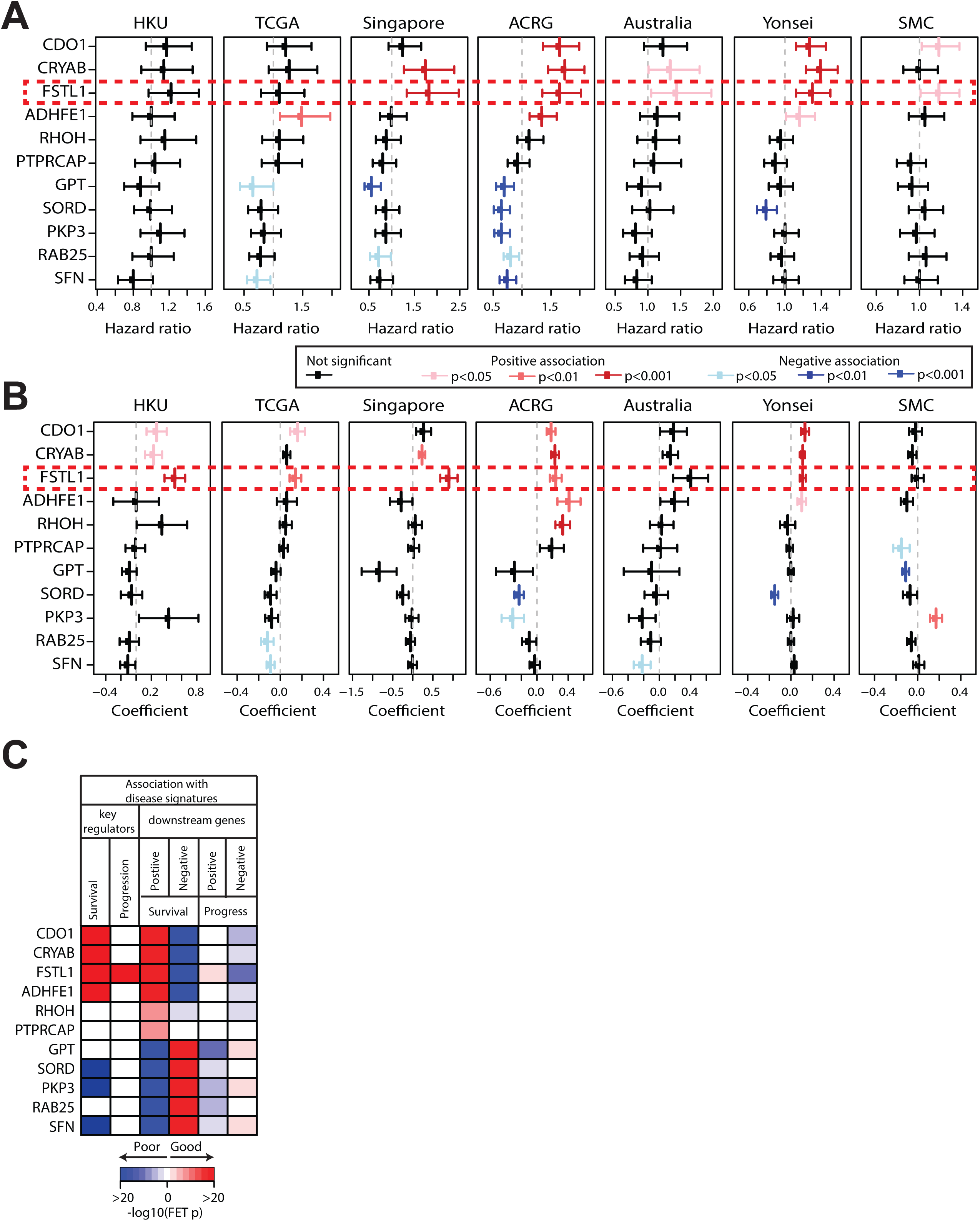
Associations between the methylation-driven key regulators and disease phenotypes. **A.** Univariate survival analysis based on expression of the 11 methylation-driven key regulators in 7 datasets. Hazard ratios with 95% confidence intervals were measured with corresponding p-value (Methods). The significant association is marked in red for poor prognosis and in blue for good prognosis. **B.** Association of expression of the 11 methylation-driven key regulators with tumor stages in 7 datasets. The coefficient of expression of each key regulator in a regression model *stage ∼ age + gender + expression* was measured with standard errors. The significant association is marked in red for advanced and in blue for early stages. **C.** The downstream genes were also compared with disease signatures such as survival-associated genes from ACRG dataset and progression signatures from Vecchi et al. (Methods). Here, the associations with signatures with good prognostics are shown in blue while the ones with bad prognostics are in red.

On the other hand, the expression levels of the G3 genes (*GPT*, *SORD*, *PKP3*, *RAB25, and SFN*) were associated with good prognosis and early stages (Figures 7A&B, Supplementary Table 6). It has been shown that loss of PKP3 protein expression indicated an invasive phonotype by comparing immunohistochemistry of PKPs in gastric tumor and normal gastric tissues^49^. Functional roles of other tumor suppressor like genes such as *GPT*, *SORD*, and *SFN* in GC have not been addressed before.

We further compared the downstream genes of the methylation driven key regulators (Supplementary Table 3) with gastric cancer survival defined based on ACRG dataset and progression signature based on early and advanced GC profiles in the Vecchi et al. study^50^ (Methods). These signatures were further split into “good” or “poor” according to the direction of their associations with patient survival (Supplementary Table 7). Most of the methylation-driven key regulators, except for *RHOH* and *PTPRCAP*, were survival-associated genes and their downstream genes were also significantly overlapped with survival signatures (FET p-value < 10^-20^, Figure 7C). The downstream genes were also consistently associated with gastric cancer progression signatures (Figure 7C). It is worth to note that downstream genes of *FSTL1* showed the strongest association with both survival and progress signatures among all methylation-driven key regulators, which was consistent with our observations that *FSTL1* expression was significantly associated with poor survival and tumor stages in multiple GC datasets (Figures 7A&B, Supplementary Table 6).

### Tumor intrinsic variations of the methylation-driven key regulators in cancer cells

Tumor-stroma interactions in GC are associated with prognosis^30–32^ and contribute to molecular heterogeneity^21^. Hence, we investigated whether variations of the methylation-driven key regulators were from tumor cells or associated with TME. First, we checked the methylation and expression levels of the 11 methylation-driven key regulators in CCLE gastric cancer cell lines^51, 52^. Based on gene expression of CCLE cancer cell lines, *FSTL1* was the only gene in the G1 group having largely varying expression among 36 cell lines while others such as *CDO1*, *CRYAB*, and *ADHFE1* were not highly expressed (Figure 8A). Interestingly, *FSTL1* expression increased in tumor tissues compared to non-tumor gastric tissues consistently in TCGA and HKU datasets while the other three showed an opposite pattern (Figure EV2). Considering their expressions had highly correlated each other (Figure 4A), *FSTL1* expression might be derived more from tumor cell variations than *CDO1*, *CRYAB*, and *ADHFE1*. Especially, *CDO1* was low expressed in cancer cells (log2(RSEM)<1 in 34 out 36 cell lines) suggesting pro-tumor progression property of *CDO1* might be not from tumor cells but other factors. Among the genes in the G3 group (*GPT, SORD, PKP3, RAB25, and SFN*), which tumor-suppressor like properties were observed in, *GPT* was less expressed than others (Figure 8A) as well as down-regulated expression in tumor tissues from non-tumor gastric tissues (Figure EV2). These results suggest that the source of expression variations of the methylation-driven key regulators was not the same even though their expressions were highly correlated in bulk tissues.

**Figure 8.**
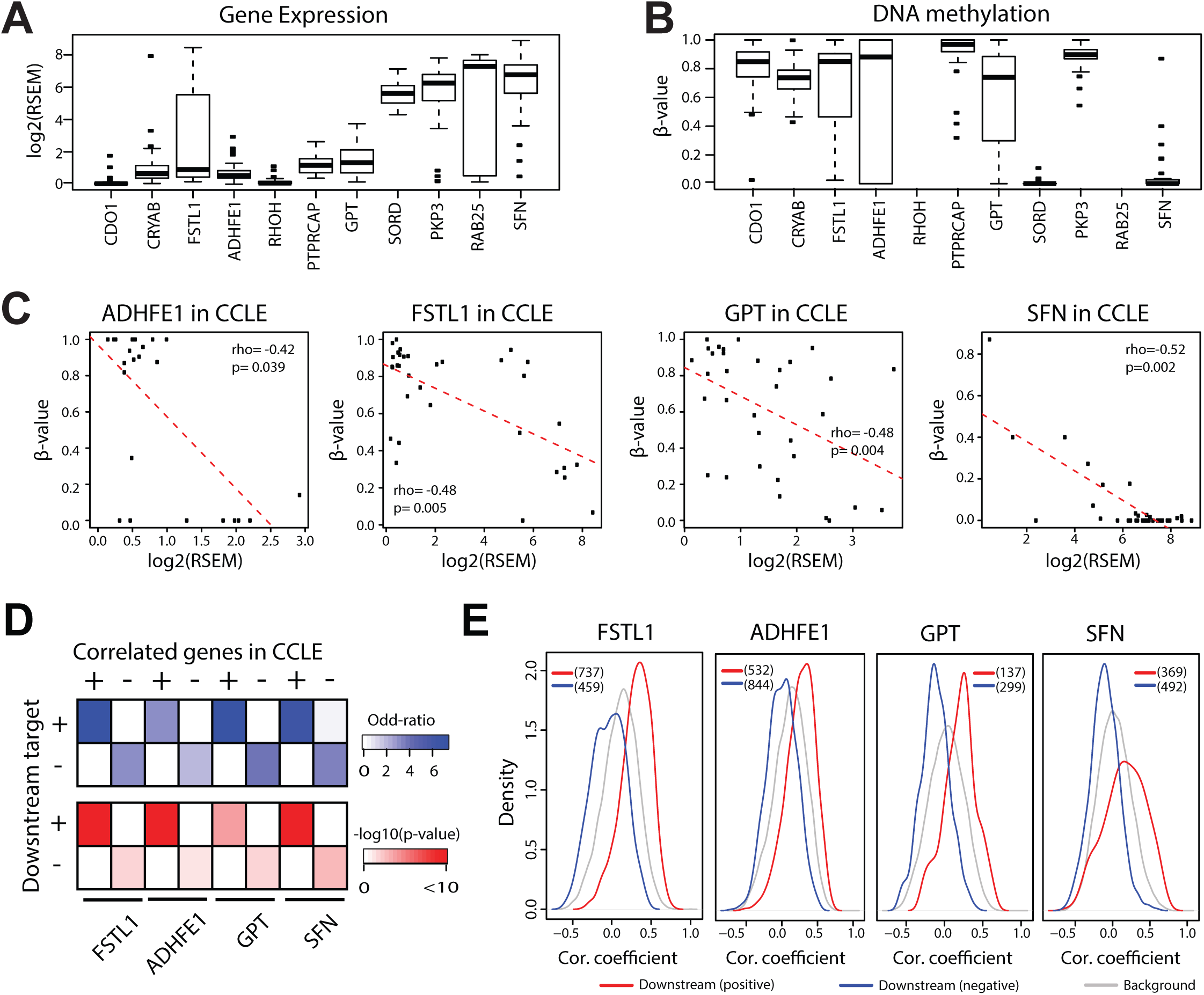
Tumor intrinsic variations associated with the methylation-driven key regulators. **A.** Distribution of gene expression (log2(RSEM)) of the methylation-driven key regulators in 36 CCLE gastric cancer cells. **B.** Distribution of DNA methylation level (β-value) within 1kb from TSS of the 11 methylation-driven key regulators in 33 CCLE gastric cancer cells. β-values for *RHOH* and *RAB25* were not available. **C.** Four methylation-driven key regulators (*ADHFE1*, *FSTL1*, *GPT*, and *SFN*) showing significant correlation between promoter methylation and gene expression in 33 CCLE gastric cancer cells. Spearman correlation coefficients (rho) with p-values are shown. **D.** Association between downstream genes identified based on ISCT and co-expressed genes for 4 methylation-driven key regulators (*FSTL1, ADHFE1, GPT, and SFN*) in CCLE gastric cancer cells. The association strengths were measured by Odd Ratios and p-value (-log10) from FET using common genes between CCLE and our three primary datasets for ISCT as gene universe (N=14581). **E.** For the 4 methylation-driven key regulators, the correlation coefficients of their downstream genes in CCLE cells were compared with those of the background genes. The red indicates density for positively regulated downstream and the blue for negatively regulated downstream genes. Gray indicates distribution of correlation coefficients of background genes. The number indicates the number of downstream genes for each key regulator covered in CCLE RNAseq data.

The CCLE methylation data within 1kb from TSS for the methylation-driven key regulators showed opposite patterns from the gene expression (Figure 8B). Four methylation-driven key regulators (*ADHFE1*, *FSTL1, GPT, and SFN*) were anti-correlated between the methylation level and gene expression (Figure 8C) suggesting that these genes were suppressed by promoter methylation in gastric cancer cells.

Furthermore, to test whether the transcription of downstream genes was driven by expression variations of the methylation-driven key regulators in gastric cancer cells, we compared the gene expression of the methylation-driven key regulators and their downstream genes in GC cell lines in two different ways. First, we identified significantly correlated genes with the 4 methylation-driven key regulators (*FSTL1*, *ADHFE1*, *GPT*, and *SFN*), which were *cis*-regulated by their promoter methylation level in GC cell lines (Figure 8C), then compared the downstream genes based on bulk tissue data and correlated genes based on cell line data. The downstream genes of the all of the 4 methylation-driven key regulators significantly enriched for genes significantly correlated with the corresponding key regulators in the CCLE GC cell lines and also the directions of the associations were consistent for all 4 genes (Figure 8D). Next, we also compared the distribution of the tumor intrinsic correlation between the methylation-driven key regulators with their downstream genes with that of background genes. The downstream genes showed clear different associations with their key regulators compared to background genes with consistent direction of correlation (Figure 8E). The cumulative distributions of the correlation coefficients were compared between the downstream genes and background genes via Kolmogorov-Smirnov (KS) Tests and all comparisons were statistical significant (KS-test D=0.27 and 0.39 with p-values=3.7×10^-30^ and 7.4×10^-96^ for positively and negatively regulated downstream genes of *FSTL1*, Supplementary Figure 5).

These results suggest that the expression of the methylation-driven key regulators was *cis*-regulated by their promoter region methylation and the putative causal relationships with their downstream genes were partially driven by tumor intrinsic variations of the genes.

### Influence of TME on the methylation-driven key regulators

Next, we examined cellular compositions of the bulk gastric tumors based on gene expression and DNA methylation (MethylCIBERSORT)^53^ (Methods). Cell proportions based on gene expression and DNA methylation profiles were similar (Pearson correlation p=2.3×10^-37^, 3.9×10^-58^, and 6.4×10^-48^ for immune, stromal, and cancer cells, respectively, Figure EV3). The expression levels of the 11 methylation-driven key regulators were significantly correlated with cell compositions (Supplementary Figure 6). The genes in the G1 group (*CDO1, CRYAB, FSTL1, and ADHFE1*) were highly associated with stromal proportions while ones in the G2 group (*RHOH and PTPRCAP*) were significantly correlated with immune proportions. The expression levels of the G3 genes (*GPT, SORD, PKP3, RAB25, and SFN*) were anti-correlated with stroma and immune proportions.

Next, we investigated the distribution of cellular compositions in the tumor clusters. In general, the immune and stromal cell proportions increased from the C1 to the C3 clusters while cancer cell proportions decreased (Supplementary Figure 7). Furthermore, the cellular compositions were tested for their association with patients’ survival (Supplementary Table 8). As the stromal proportions were correlated with the expression of the methylation-driven key regulators in the G1 group, they were significantly associated with poor survival in multiple datasets (Hazard Ratios (HRs)= 1.41, 1.62, 1.35 and 1.22, p-values= 0.008, 5.4×10^-8^, 2.4×10^-6^ and 0.02 for Singapore, ACRG, Yonsei and SMC, respectively) consistently with previous reports showing poor survival associated with higher tumor-stroma interactions^30–32^.

As the methylation-driven key regulators potentially regulated a large number of downstream genes in each dataset (Figure 3D) and the methylation-driven key regulators were highly correlated with cell type proportions, we also tested whether the downstream genes were influenced by cell type proportion as well. Indeed, downstream target genes showed stronger association with cell type proportions compared to other *trans* genes suggesting influence of TME in the causality test (Appendix and Supplementary Figure 8). The results based on the CCLE cancer cell lines as well as cell type proportions together suggest that the transcription regulations of the methylation-driven key regulators on their downstream genes were influenced by both TME and intrinsic variations of cancer cells.

Taken all results together, tumor cells and their interactions with TME may drive molecular and cellular heterogeneity of gastric cancer which, in turn, leads to heterogeneity of clinical outcomes of GC patients.

## Discussions

Gastric cancer is of highly heterogeneous molecular and histological features. These molecular and cellular heterogeneities were driven by complex and diverse genomic and epigenetic alterations occurred during tumorigenesis and progression of gastric cancer. We integrated DNA methylation, CNV, and gene expression data to understand these heterogeneities, and developed the ISCT procedure to infer causal relationships between a pair of *cis*-methylation gene and *trans* genes. Among the numerous numbers of significantly associated *cis*-*trans* gene pairs, ISCT identified potential causal gene pairs whose significant associations were mediated by expression of *cis* genes that were driven by methylation variations. Based on simulation studies, we showed that ISCT outperformed mediation tests with consistently higher detection power and was robust against to colinearity problems. It is not trivial to systematically evaluate the performance of ISCT in empirical data, thus we focused only on coherent common observations from the three multi-omics GC datasets (HKU, TCGA, and Singapore). In addition, we collected gene expression data from multiple independent GC cohorts to cross-validate our observations.

By applying ISCT, we identified 11 methylation-driven key regulators common from all three datasets. These genes were further grouped into three groups based on their gene expression similarity (Figure 4A): G1 (*CDO1, CRYAB, FSTL1, and ADHFE1*), G2 (*RHOH and PTPRCAP*), and G3 (*GPT, PKP3, RAB25, SFN, and SORD*). The G1 genes showed strong association with poor survival while the expression of G3 was associated with good prognosis (Figure 7).

Based on expression of the 11 methylation-driven key regulators, GC samples were grouped into three clusters (C1 to C3) which showed distinct molecular or clinical features (Figure 4). Samples in C3 cluster were significantly enriched for diffuse type tumors in all datasets while samples in C1 were for intestinal type ones. In addition, C3 samples were enriched for the GS subtype defined in TCGA cohort. While MSI tumors were well known to have CIMP features^54^, there was no previously reported association between methylation with GS tumors. The C3 samples were also enriched for mesenchymal-types (or invasive) features determined in Singapore, ACRG, Australia, and CPTAC cohorts. Interestingly, we observed correlation of the sample clusters and tumor stage, and samples in C3 showed enrichment for advanced stages and also worst survival outcomes in multiple cohorts (HKU, TCGA, Singapore, ACRG, Yonsei and SMC dataset, Figure 4D). Previous studies about DNA methylation in GC mostly focused on molecular subtypes defined based on CIMP status^9, 12, 55^ but the association of CIMP tumors with clinicopathological features were not well agreed in all studies. ISCT enables us to identify methylation-driven key regulators. The associations of the key methylation-driven regulators with molecular and clinical features were consistent across multiple independent GC cohorts (Supplementary Tables 4 and 6).

Comparing tumor clusters based on methylation-driven key regulators with reports by Oh et al., the C1 and C3 tumors were enriched for EP and MP, respectively. Moreover, we showed the adjuvant chemotherapy sensitivity in C1-3 or EP/MP clusters was stage dependent, suggesting significance of considering not only molecular features but also progression of tumors when treating GC patients.

Some of the methylation-driven key regulators were reported to play a significant role in GC. Chen et al. showed that overexpression of *CRYAB* induced EMT, migration, and invasion of gastric cancer cells in *vitro* and in *vivo* as well as reversing these phenomena by silencing *CRYAB*^47^. In addition, they showed strong associations between *CRYAB* expression and cancer metastasis and survival outcomes in patients. These results were consistent with our results suggesting *CRYAB* as a potential oncogene regulating EMT associated genes (Figure 7 and Figure EV1B). Demirag et al. investigated IHC of plakophilins in gastric adenocarcinoma and normal gastric tissues and reported low PKP3 protein levels were correlated to the node number, tumor stages, and poor prognosis in gastric carcinoma^49^, which was consistent with our result that *PKP3* was associated with good survival and down-regulated in advanced stages. *ADHFE1* was reported as an oncogene by inducing metabolic reprogramming in breast cancer^56^, but no previous association with gastric cancer was reported. Interestingly, Oh et al. also observed upregulation of *IGF1* coupled with promoter region hypomethylation in mesenchymal gastric tumors and showed inhibition of *IGF1* reduced tumor growth in mesenchymal type tumor cells^46^. *IGF1* was identified as one of methylation-driven key regulators in the Singapore and TCGA dataset (Supplementary Table 2).

Contrary to those positive controls, the observation with Cysteine dioxygenase type 1 (*CDO1*) was not consistent with previous reports. Hao et al. showed that suppressing CDO1 increased ferroptosis resistance in human gastric cancer cells and tumors in CDO1 knockdown mice grew faster compared to controls^57^. Harada et al. reported hypermethylation of *CDO1* promoter as an independent prognostic marker in gastric cancer^58^. This discrepancy could be explained by very low expression in cancer cells (Figure 6A), downregulated expression in tumors compared to normal tissues (Figure EV2), and strong association of *CDO1* expression with stromal proportions (Supplementary Figure 6), together suggesting that the *CDO1* expression in bulk tumors was mainly determined by stroma cell proportion, which is an independent prognostic factor in GC^30^.

Other methylation-driven key regulators have not been previously associated with gastric cancer. Follistatin like 1 (*FSTL1*) was associated with tumor cell proliferation, migration, and invasion in several other cancers including lung, colon, breast and renal cell carcinoma^59–62^, but had not distinctively been linked with gastric cancer. Our results suggest *FSTL1* as a potential novel oncogene in gastric cancer based on its over-expression in tumor samples and strong association with patients’ survival and tumor stages in multiple GC datasets (Figure 7, Supplementary Table 6). While *FSTL1* expression in the bulk was also strongly correlated with stromal proportions like *CDO1* (Supplementary Figure 6), *FSTL1* expression was associated with its promoter methylation in gastric cancer cells (Figure 8A-C). Moreover, *FSTL1* and its downstream genes were significantly correlated within cancer cells (Figure 8D&E). These observations suggest that *FSTL1* expression is governed by both tumor intrinsic variation as well as TME. On the other hand, Glutamic-Pyruvic Transaminase (GPT) showed exact opposite patterns from *FSTL1*. The *GPT* expression was suppressed by hypermethylation in tumors compared to normal tissues and was associated with good survival, suggesting its role as a tumor suppressor gene (Figure 7, Supplementary Table 6). GPT is known to play an important role in the intermediary metabolism of glucose and amino acids but was reported to be associated liver diseases^63^. While there were no literatures about *GPT* in gastric cancer, its expression was associated with good survival in liver and rectal cancer (https://www.proteinatlas.org/ENSG00000167701-GPT).

*FSTL1* and *GPT* expression levels were regulated by its promoter methylation in gastric cancer cells (Figure 8C). Even though the expression levels of the methylation-driven key regulators in bulk tumors were significantly associated with immune or stromal proportions in TME, genes correlated with methylation-driven key regulators in GC cell lines significantly overlapped with their downstream target genes inferred using ISCT from bulk tissue data (Figures 8D&E). These results suggest that tumor-TME interactions contribute to expression variations of methylation-driven key regulators such as *FSTL1* and *GPT*, which in turn give rise to the molecular and histological heterogeneity of gastric cancer. Further investigation in co-cultured system might elucidate detail roles of these genes in tumor-stroma interaction.

While our main interests are on methylation-driven key regulators, CNV alterations also play important roles in tumorigenesis and progression of GC. Using ISCT, 39 common CNV-driven key regulators were identified (details in Appendix, Supplementary Figures 9 and 10). Most of these key CNV regulators were located in chromosomes 20 and 8 where the gain of DNA copy in these locations was known in several of previous GC studies^19, 64–66^. These genes located in the chromosome 20 were significantly amplified specifically for CIN tumors, in which no clear methylation features were associated. Indeed, no CNV-driven key regulator overlapped with the methylation-driven key regulators. Interestingly, the downstream genes of the CNV-driven key regulators were shared with those of the methylation-driven key regulators (Appendix, Supplementary Figure 11). These suggest the different tumorigenic pathways through methylation or copy number alterations may have similar downstream effects.

In this study, we reported 11 genes as methylation-driven key regulators identified based on ISCT. These genes characterize diverse heterogeneities of GC showing distinct molecular, cellular, and histological features as well as clinical outcomes. They were also associated with cell type proportions suggesting their roles in TME interactions. Further investigations for their molecular functions especially *FSTL1* may reveal their novel roles in tumorigenesis and progression of GC that will enhance better diagnosis, prognosis, or treatment of GC patients.

## Materials and Methods

### GC Datasets used in integrative causal modeling

Three GC cohorts from Hong Kong University (HKU), TCGA Stomach adenocarcinoma (TCGA), and University of Singapore (Singapore), which contain gene expression, methylation, and CNV profiles, were used in this study. The HKU dataset was deposited in European Genome-phenome Archive with the study ID EGAS00001000597^8^. The molecular data for the TCGA cohort^10^ were downloaded from TCGA data portal (https://gdc.cancer.gov). The Singapore dataset was downloaded from Gene Expression Omnibus (GEO) with accession numbers GSE30601^9^, GSE15460^67^, and GSE31168^18^ for methylation, gene expression and CNVs, respectively. Prior to our integrative analysis, we excluded Epstein-Barr virus (EBV) positive samples (6 out of 98, 24 out of 235, and 5 out of 91 in HKU, TCGA, and Singapore cohorts, respectively) according to their annotation as EBV positive GC have unique and distinct DNA hyper-methylation patterns^68^. Sample alignment procedure^69^ was applied to confirm that different types of molecular data pertaining to the same individuals were matched (details in Appendix) and 92, 211, and 86 samples in the HKU, TCGA, and Singapore datasets, respectively, were finally selected for the integrative analyses. Clinical information of the three datasets is shown in Supplementary Table 1.

### Independent GC datasets for validations

Five independent cohorts with gene expression profiles were used for validating our observations based on integrative analysis: 1) Microarray profiles of 300 GC tumors from the Asian Cancer Research Group (ACRG) were downloaded from GEO with accession number GSE62254^43^; 2) A microarray dataset of 70 GC patients from Australian cohort (Australia) was downloaded from GEO with accession number GSE35809^45^; 3) A RNAseq profiling dataset described in a proteogenomic paper by Mun et al. (CPTAC) consisting of 80 patients with early onset gastric cancers was downloaded from GEO with accession number GSE122401^70^; 4) A microarray dataset from Yonsei hospital (Yonsei) consisting of 433 GC patient samples collected during 2000-2010 was downloaded from GEO with accession number GSE84437; 5) A microarray dataset consisting of 432 formalin-fixed paraffin-embedded (FFPE) tissues from Samsung Medical Center (SMC) was downloaded from GEO with accession number GSE26253^70^. For each dataset, EBV positive tumors were removed prior to following analysis. For ACRG, Australia, and CPTAC dataset, which the EBV status of the samples was available, final 221, 69, and 74 samples were selected. For Yonsei and SMC dataset without EBV status information, samples were clustered based on gene expression of the EBV signature genes^71^ and no samples were filtered out in Yonsei but 57 samples were removed in SMC dataset (details in the Appendix). The demographics of these validation datasets are also listed in Supplementary Table1.

Four more gene expression datasets from the study by Oh et al.^46^ were additionally used to investigate the association between methylation-driven key regulators and epithelial/mesenchymal phenotypes. The processed microarray data are available in GEO with accession number GSE26899 for KUGH, GSE26901 for KUCM, GSE13861 for YUSH, GSE28541 for MDACC. The EP/MP subtype for each tumor is downloaded from supplementary tables of their paper published at Nature Communication in 2018^46^.

### Data preprocessing

For the gene expression data in HKU, profiled on Illumina HT12v4, probe level data was obtained from median summarization over background corrected bead level data from Illumina Genome Studio, followed by quantile normalization on log (base 2) transformed probe intensity. Multiple probes for a gene were summarized (median) into gene level data after non-performing probes were excluded. For the RNAseq data from TCGA, RNA-Seq by Expectation-Maximization (RSEM) data downloaded and gene level expression was obtained from log transformation. For the Singapore dataset, profiled by Affymetrix UG U133A platform, probe intensity was normalized by a standard affy function, Robust Multi-array Average (RMA), with log transformation. For the five validation cohorts, we used the “getGEO” function from GEOquery package to download gene expression data as deposited in GEO database^72^.

To associate DNA methylation and gene expression, we focused on methylation variations within gene promoter regions. NCBI RefSeq annotation was downloaded in gtf format and methyl probes located within 10kb upstream from Transcription Starting Sites were selected. Since the methylation values were not normally distributed as gene expression or copy numbers^73^, beta values of each CpG probe were transformed based on rank-based normal transformation using the “rntransform” function embedded in GenABEL package (Supplementary Figure 12)^74^.

For the CNV profiles, Circular binary segmentation (CBS)^75^ of the log R ratio values was used for all three dataset. Then each segment value was mapped to gene levels based on coordinate information of RefSeq reference annotation as mapping of methyl probes. Detail information of molecular data platforms and the final number of features used in this study is summarized in Supplementary Table 9.

### An Integrative Sequential Causality Test (ISCT) for causal regulations by DNA methylation, and CNVs

Previously, we developed a causality test for modeling transcription regulations by promoter region methylations^34^. As transcriptional regulations occur at multiple levels simultaneously, here we describe a model for transcriptional regulations by methylation and CNVs in both *cis* and *trans* (Figure 1A). Given a *cis*-regulated gene *x*, (*g*_*x*_∼ *m*_*x*_ + *c*_*x*_) and a *trans*-regulated gene *y*, (*g*_*y*_ ∼ *m*_*x*_ | *m*_*y*_, *c*_*y*_), where *g*_*x*_ and *g*_*y*_ are expression levels, *c*_*x*_ and *c*_*y*_ are corresponding copy number variations, and *m*_*x*_ and *m*_*y*_ are promoter region methylation levels of gene *x* and *y*, respectively, the causal relationship between *cis* gene expression and *trans* gene expression holds when the *trans*-regulation can be completely explained by the *cis* gene expression. In other words, we hypothesize that the *trans* relationship (*g*_*y*_ ∼ *m*_*x*_ | *m*_*y*_, *c*_*y*_) arises from the probability chain *p*(*m*_*x*_ → *g*_*x*_ → *g*_*y*_|*c*_*x*_, *m*_*y*_, *c*_*y*_), which can be decomposed as a production of probabilities of a chain of statistical tests as below. The causal relationship is significant when the *trans* relationship (*g*_*y*_ ∼ *m*_*x*_ | *m*_*y*_, *c*_*y*_) becomes non-significant after conditioning on the *cis*-gene expression *g*_*x*_, in which case the *trans* relationship between *m*_*x*_ and *g*_*y*_ is “caused” by *g*_*x*_.

Similar to the mediation test outlined by Baron and Kenny^36^, the causality test can be broken down into steps: (1) the *cis* regulation: can be modeled as a linear regression *g*_*x*_ ∼ *m*_*x*_ + *c*_*x*_; (2) the *trans* association between *g*_*y*_ and *m*_*x*_: instead of being modeled as a linear regression *g*_*y*_ ∼ *m*_*x*_ + *m*_*y*_ + *c*_*y*_, was modeled in a sequential process: we accounted *cis* regulations *g*_*y*_∼ *m*_*y*_ + *c*_*y*_ and identified residual variance that could not be explained by *cis* regulation 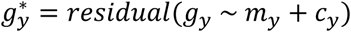, then we modeled the *trans* association as 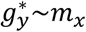; (3) the association between *g*_*x*_ and *g*_y_: was modeled similarly as 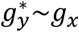; (4) the conditional independence (the indirect effect) between *m*_*x*_ and *g*_*y*_|*m*_*y*_, *c*_*y*_ was modeled in a sequential procedure: identifying the residual variance that could not be explained by *g*_*x*_ as 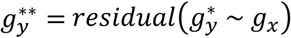 then the conditional independence was assessed as 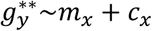 instead of a standard linear regression *g*_*y*_∼*m*_*x*_ + *c*_*x*_ + *g*_*x*_ + *m*_*y*_ + *c*_*y*_. As we previously described causal relationship between promoter region methylation and trans gene expression^34^, *p*(*m*_*x*_ → *g*_*x*_ → *g*_*y*_|*c*_*x*_, *m*_*y*_, *c*_*y*_) was mainly determined by *p*(*m*_*x*_ ⊥ *g*_*y*_|*g*_*x*_, *m*_*y*,_*c*_*y*_) given significant *cis* and *trans* relationships for gene *x* and *y* (FDR<0.05).

Similarly, the causal relationship between a *cis* CNV gene*x* and a *trans* gene*y* was modeled as *p*(*c*_*x*_ → *g*_*x*_ → *g*_*y*_|*m*_*x*_, *m*_*y*_, *c*_*y*_).

### Comparison with mediation tests

A mediation test is an alternative to the ISCT for testing causal relationships between the *cis* gene methylation and *trans* gene expression. In a mediation model, the *cis* methylation *m*_*x*_ can be perceived as the independent variable, the *trans* gene expression *g*_*y*_ is the dependent variable, and the *cis* gene expression *g*_*x*_ is the potential mediator. Various mediation test methods exist for testing whether the relationship between the independent variable and the dependent variable is mediated through the potential mediator. One of the most widely used approaches to test for mediation is the causal steps method^35, 36, 76^, which evaluates three regression models: the first one assesses whether the independent variable affects the mediator by regressing the mediator on the independent variable; the second one assesses whether the independent variable affects the dependent variable by regressing the dependent variable on the independent variable; the third one assesses whether the mediator affects the dependent variable when the independent variable is controlled by regressing the dependent variable on both the independent variable and the mediator. The mediation effect is established if all three regressions show significant relationships, and the effect of the independent variable on the dependent variable is reduced in its absolute size after controlling for the mediator. Moreover, if the independent variable shows no effect on the dependent variable in the third regression, the mediation is full. Another method to test for mediation is the Sobel test,^37^ which evaluates the significance of the mediation (indirect) effect by comparing its magnitude divided by its estiamted standard error of measurement to a normal distribution. The Sobel test is known to be conservative^77^ because of its normal approximation of the test statistic. Nevertheless, both mediation tests are expected to be underpowered in testing whether the association between the independent variable *m*_*x*_ and the dependent variable *g*_*y*_ is (completely) mediated through the mediator *g*_*x*_, given that *cis*-regulation indicates colinearity between the mediateor *g*_*x*_ and the independent variable *m*_*x*_. The ISCT approach, on the other hand, assigns variances to each variable according to the sequence of biological events so that it does not suffer from the colinearity problem. For example, in an extreme case where *g*_*x*_ and *m*_*x*_ are perfectly correlated, both coefficients will be non-significant in the regression 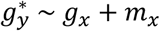, giving non-significant result for the mediation test; whereas given significant trans-regulation between 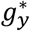 and *m_x_*, the first regression in the ISCT approach 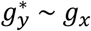 will give significant coefficient for *g_x_*, and the second regression 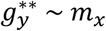 will give a non-significant coefficient for *m*_*x*_, resulting in a significant case for the sequential regression approach. In reality, most of the *g*_*x*_ and *m*_*x*_ tested for mediation or causality are not perfectly correlated but may be highly correlated as a significant *cis*-regulation for the pair is required. Therefore, the mediation test is expected to give overly conservative results because of its reduced power in the presence of colinearity, while the ISCT approach is expected to be better powered in detecting the causal relationships.

### Simulations for comparing ISCT and mediation tests

To estimate the power and the false postive rate of each method to detect the underlying causal/mediation relationships, we conducted two simulation studies based on a causal model and an independent model.

#### 1) Simulation #1

We randomly selected 10,000 *causal* pairs from the HKU cohort with significant *cis*- and *trans*-relationships that were tested significant by ISCT. We preserved the values of the independent variable *m*_*x*_ and the potential mediator *g*_*x*_ (and the covariates *c*_*x*_, *m*_*y*_), and simulated 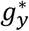 based on the mediator *g*_*x*_ so that 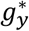 is correlated with *g*_*x*_ at the same correlation level between *g*_*x*_ and the original 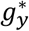 values in the HKU cohort, then *g*_y_ is calculated from the simulated 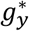 and its original regression coefficient. This data-generating process mimiced the effect of a mediator^38^, by leaving the correlation between the dependent variable and the independent variable *cor*(*g*_*y*_, *m*_*x*_) to vary solely as a result of the causal path from *m*_*x*_ → *g*_*x*_ → *g*_*y*_. We selected simulated *g*_*y*_ values showing significant *trans*-relationship with *m*_*x*_, then applied both ISCT and two mediation tests to detect the underlying causal/mediation relationships.

#### 2) Simulation #2

We randomly selected 10,000 *non-causal* pairs from the HKU cohort with significant *cis*- and *trans*-relationships that were tested non-significant by ISCT. We preserved the values of the independent variable *m*_*x*_ and the potential mediator *g*_*x*_ (and the covariate *c*_*x*_), and simulated *m*_*y*_ based on *m*_*x*_ at a correlation level sampled from the correlation distribution between methylation levels of a meth-probe and that of its trans-associated gene’s most associated meth-probe in the HKU cohort, then simulated the dependent variable *g*_*y*_ based on *m*_*y*_ at a correlation level sampled from the correlation distribution between all genes and their most associated meth-probes in the HKU cohort. This data-generating process mimiced a trans relationship resulting from a path independent of *g*_*x*_ : *m*_*x*_ → *m*_*y*_ → *g*_*y*_. We selected simulated *g*_*y*_ values showing significant *trans*-relationship with *m*_*x*_, then applied both ISCT and two mediation tests to detect the underlying causal/mediation relationships.

Furthermore, we performed two additional simulation studies given certain correlation levels among the variables to investigate the performance of each method under each specific scenario in the presence of colinearity.

#### 3) Simulation #3

We randomly selected 10,000 *cis*-*trans* pairs from the HKU cohort with significant *cis*- and *trans*-relationships. We preserved the values of the independent variable *m*_*x*_ (and the covariate *c*_*x*_) and the potential mediator *g*_*x*_, and generated the dependent variable 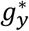 so that 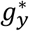 is correlated with the mediator *g*_*x*_ at given correlation levels. Then we selected *cis*-*trans* pairs and applied ISCT and two mediation tests to detect the underlying causal/mediation relationships.

#### 4) Simulation #4

We randomly selected 1,000 *cis*-*trans* pairs from the HKU cohort, and preserved the values of the independent variable *m*_*x*_ (and the covariate *c*_*x*_), and simulated the potential mediator *g*_*x*_ so that *g*_*x*_ correlated with the independent variable *m*_*x*_ at a pre-specified correlation level *cor*(*g*_*x*_, *m*_*x*_), then simulated the dependent variable so that the correlation between the dependent variable and the mediator at a pre-specified level. Then we selected *cis*-*trans* pairs and applied ISCT and two mediation tests to detect the underlying causal/mediation relationships.

### Key regulator identification

Key regulators were determined based on the number of downstream genes. For methylation-driven key regulators, causal relationships between *cis* and *trans* genes were assessed based on individual methylation probes, summarized at gene levels as there were multiple methylation probes profiled in the promoter regions of each individual gene. *Cis* methylation and CNV genes were sorted based on the number of their downstream genes. Then key regulators were defined as ones whose numbers of downstream genes were significantly higher compared to others. The cutoff for defining key regulators in each dataset was set based on the reflection point from numbers of downstream genes for each regulator^78, 79^.

### GC related signatures

GC progress signatures were defined as up and down-regulated genes in advanced GCs compared to early stage ones^50^. GC survival associated genes were derived based on Asian Cancer Research Group (ACRG) GC cohort^43^, which includes only gene expression^67^. Only samples with living without recurrence or samples with death due to disease were used to define survival signatures. The association of expression of each gene with survival information was tested using a Cox regression model as *survival ∼ age + gender + expression*. In total, 3375 GC survival associated genes were identified as FDR<0.01.

### Functional analysis

To identify enriched function among a set of selected genes, a collection of Hallmark gene sets, curated gene sets, and GO terms in Molecular Signatures Database (MSigDB) were used^42^. The significance of the overlap with query genes was tested via the Fisher’s exact test (FET).

### Survival analysis

Clinical information for samples in all datasets was downloaded from their corresponding papers or GEO database. Cancer specific survival (CSS) was available for HKU, TCGA, and ACRG datasets and recurrence free survival was used for SMC dataset. For Singapore, Australia, and Yonsei datasets, overall survival (OS) was used. CPTAC dataset was omited for survival analysis because the events occurred in only 9 samples out of 74 samples, so that it was not sufficient to perform survival analysis in the CPTAC dataset. For univariate survival analysis with age and gender as covariates was used as *survival ∼ age + gender + factor*, where factors were gene expression, cell type proportions, or clusters. R package “survival” was used for the survival analysis.

### Cell component decomposition

CIBERSORT^80^ (https://cibersort.stanford.edu/) was used to decompose cell components into immune, stromal and cancer proportions. For the immune references, the original LM22 data was used and proportions of individual 22 immune cell types were summed up to immune proportions. For stroma and cancer cells references, we downloaded microarray CEL files (Affymetrix HG_U133+2) of 6 stomach fibroblasts (3 submucosal and 3 subperitoneal fibroblasts) from GSE63626^81^ and 36 stomach carcinoma cells from Cancer Cell Line Cyclopedia (CCLE)^51^ (https://portals.broadinstitute.org/ccle). The cell profiles were processed to generate a signature matrix by comparing one cell type versus all other cell types. And the signature matrix was used to the proportions of immune, fibroblast, and cancer cells in samples of the 8 GC cohorts (3 primary and 5 validation datasets). Cell type proportions based on DNA methylation were downloaded from MethylCIBERSORT^53^ results page (https://zenodo.org/record/3242689#.XQ0S9vlKjOR).

## Expanded View Figure legends

**Figure EV1.**
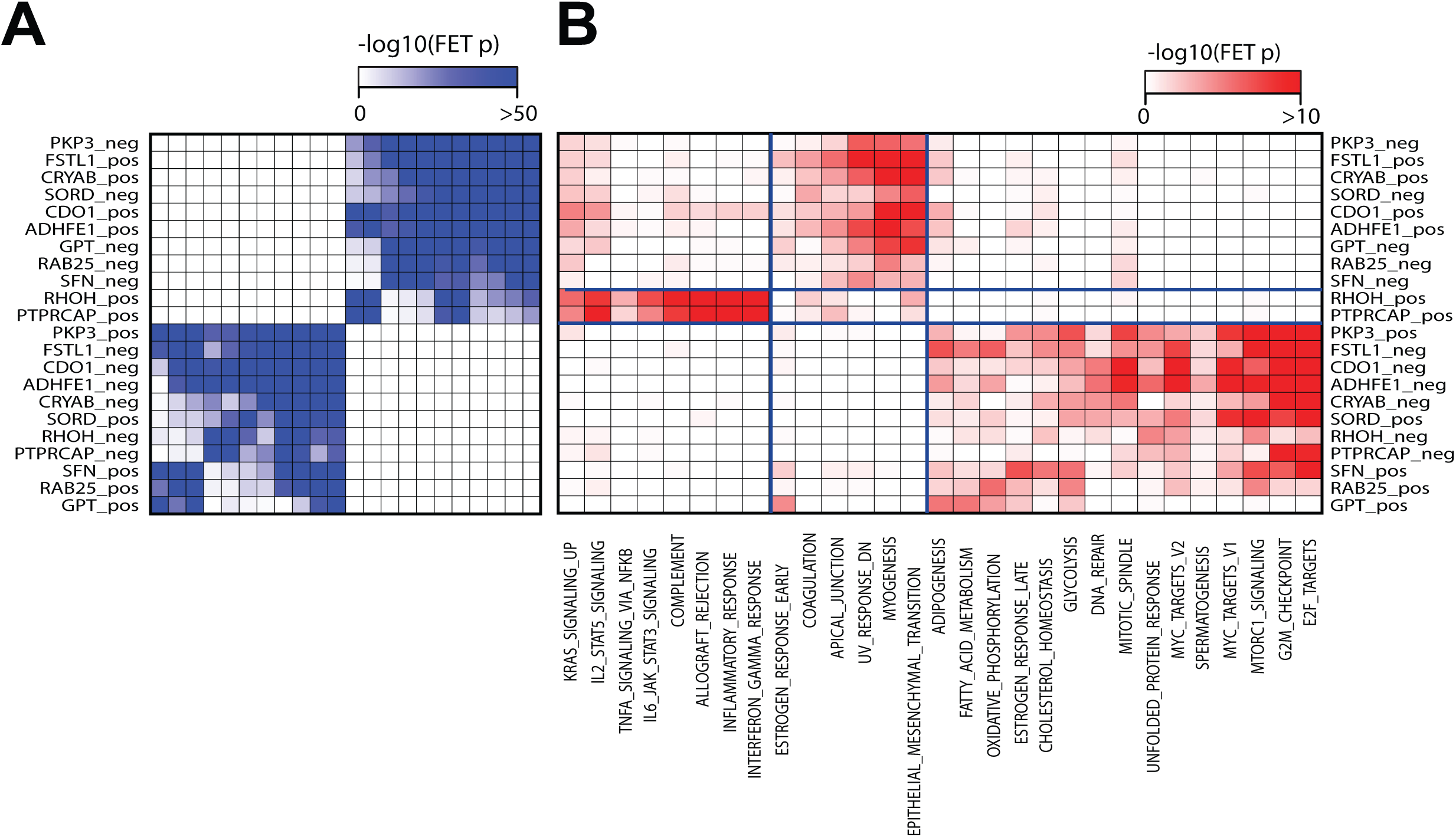
Downstream target genes of the methylation-driven key regulators. A. The overlap of downstream genes of the 11 methylation-driven key regulators. The significance of the overlap was measured by FET p-value (-log10(p-value)). B. The downstream genes were compared with MSigDB Hallmark gene sets and significantly associated gene sets (p<0.001 from multiple testing) with any downstream gene sets are shown.

**Figure EV2.**
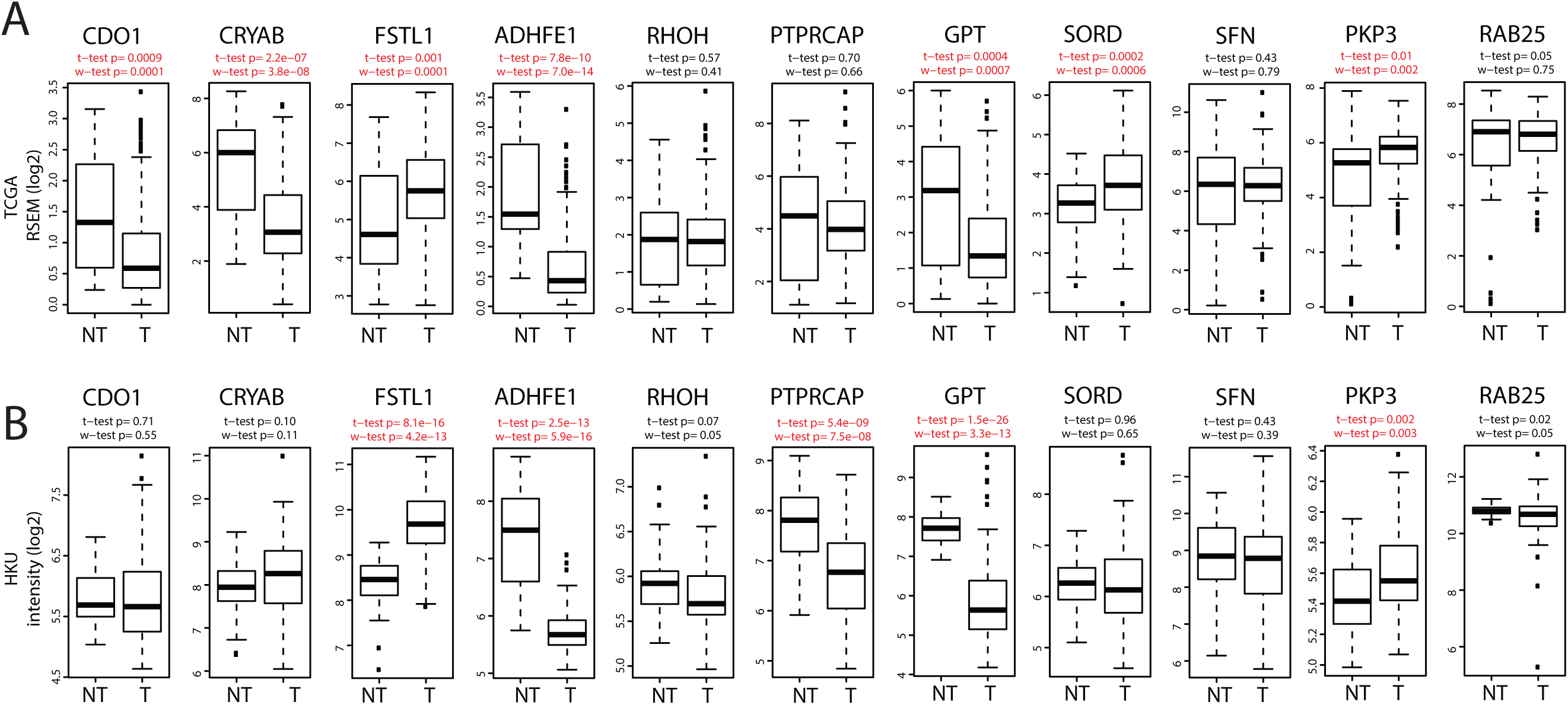
Gene expression difference of the methylation-driven key regulators in tumors compared to normal tissues. The expression levels of the methylation-driven key regulators were compared between normal and tumors based on Student t-test and Wilcox Rank Sum test A. in TCGA (211 tumors vs. 27 normal tissues) and B. in HKU (92 tumor vs. 35 normal tissues). The significant differences (p<0.01) are marked in red. Normal tissues are not available for Singapore dataset.

**Figure EV3.**
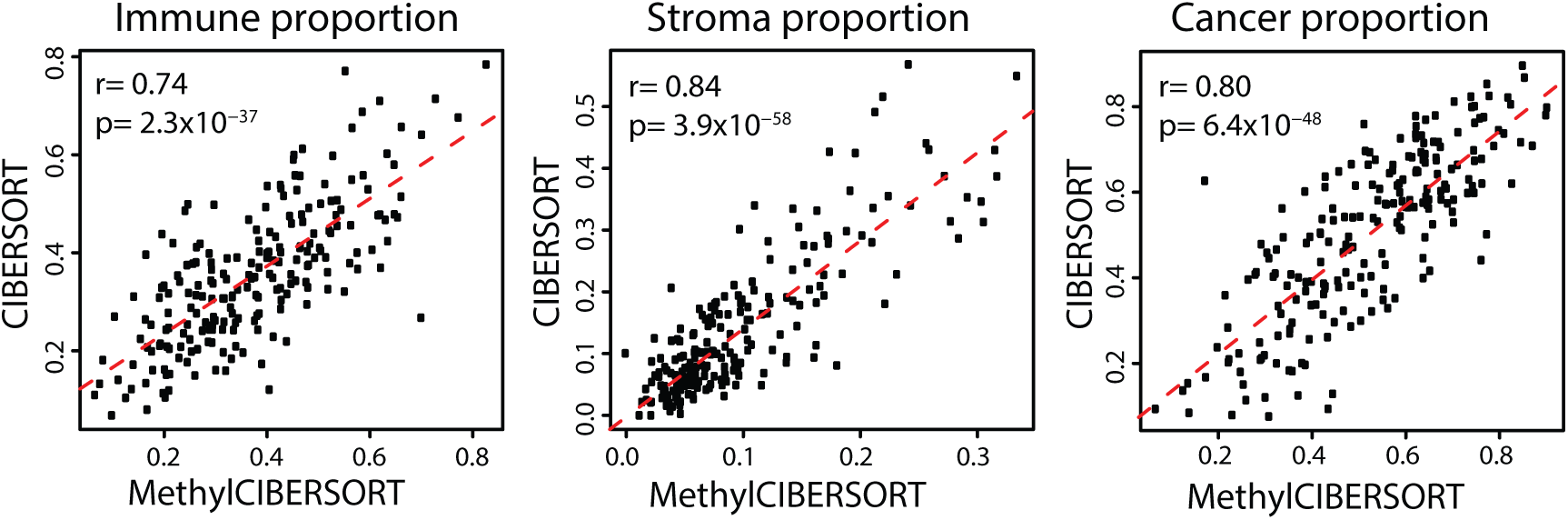
Comparison of cell types proportions based on DNA methylation (MethylCIBERSORT) and gene expression (CIBERSORT). The cell type proportions of 211 TCGA samples were measured based on DNA methylation and gene expression were compared. Pearson correlation and corresponding p-values were measured for immune, stromal, and cancer proportions.

